# Stable behavioral state-specific large scale activity patterns in the developing cortex of neonates

**DOI:** 10.1101/2021.02.19.431943

**Authors:** Nima Mojtahedi, Yury Kovalchuk, Alexander Böttcher, Olga Garaschuk

## Abstract

Endogenous neuronal activity is a hallmark of the developing brain. In rodents, a handful of such activities were described in different cortical areas but the unifying macroscopic perspective is still lacking. Here we combined large-scale *in vivo* Ca^2+^ imaging of the dorsal cortex in non-anesthetized neonatal mice with advanced mathematical analyses to reveal unique behavioral state-specific maps of endogenous activity. These maps were remarkably stable over time within and across experiments and used patches of correlated activity with little hemispheric symmetry as well as stationary and propagating waves as building blocks. Importantly, the maps recorded during motion and rest were almost inverse, with sensory-motor areas active during motion and posterior-lateral areas active at rest. The retrosplenial cortex engaged in both resting- and motion-related activities, building functional long-range connections with respective cortical areas. The data obtained bind different region-specific activity patterns described so far into a single consistent picture and set the stage for future inactivation studies, probing the exact function of this complex activity pattern for cortical wiring in neonates.

## Introduction

Mammalian cortex represents a complex computational space with information processing units interconnected via local and long-range connections. The initial formation of cortical architecture is laid out by molecular cues but its refinement critically depends on neuronal activity (Katz and Shatz, 1996; Kirkby et al., 2013; Luhmann et al., 2016). In the rodent forebrain, many of the initial coarse connections are refined during the first postnatal week leading, for example, to the eye-specific input segregation in the lateral geniculate nucleus of the thalamus and formation of the retinotopic map in the visual (Ackman and Crair, 2014; Kirkby et al., 2013) or the emergence of the barrel map in the somatosensory (Luhmann et al., 2016; Petersen, 2007) cortex. Because during this time period activation of these cortices through extrinsic (sensory) stimuli is rather limited (see Fig. 2 in (Hanganu-Opatz, 2010)), this refinement likely relies on intrinsic neuronal activity.

**Fig. 2.**
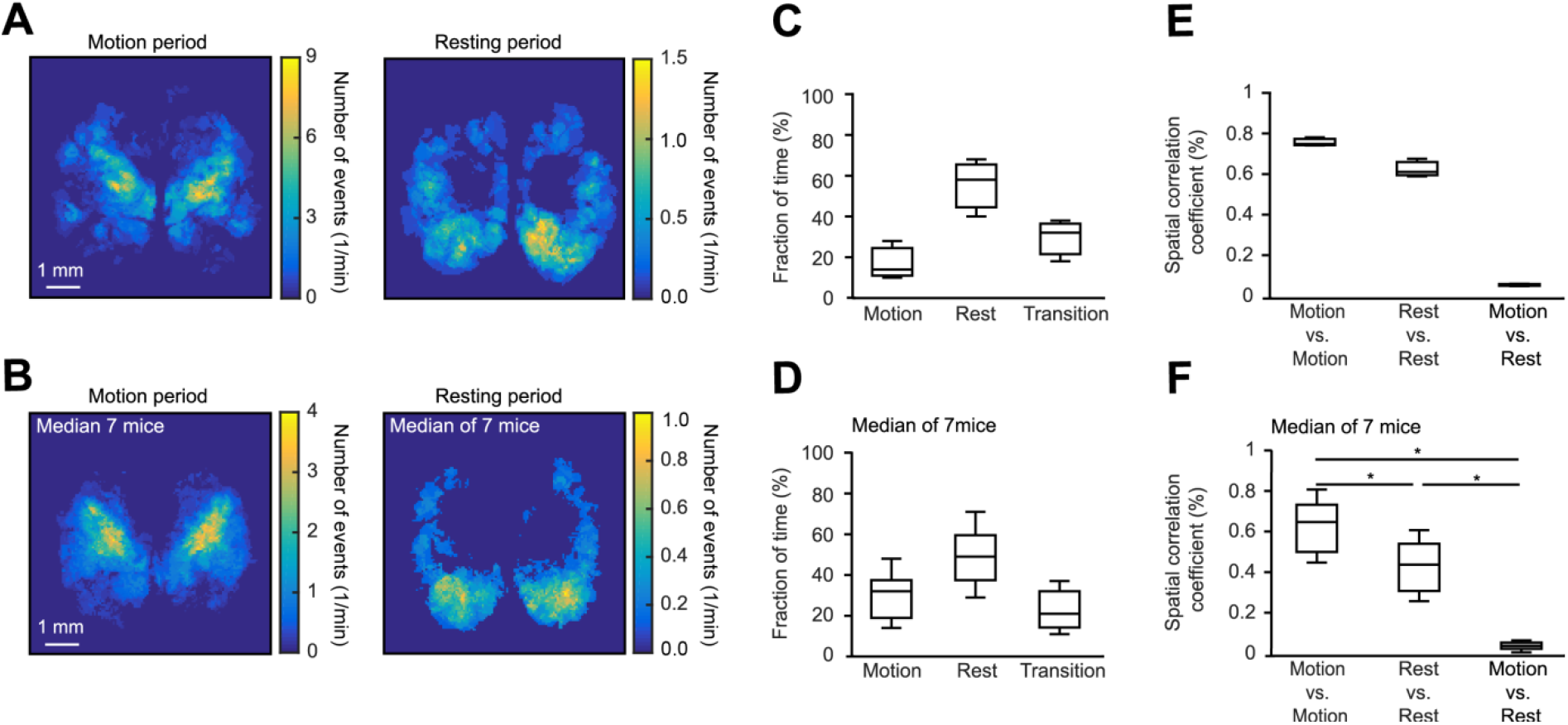
ROI-based frequency maps of local activity. ***A***, Representative frequency maps (NMF-based ROI analysis) recorded in a mouse 5 (Fig. 2-2) during motion (left panel) and resting (right panel) time periods. ***B***, Same analyses as in ***A***, but each pixel is a median of data obtained in 7 different animals. ***C***, Box-and-whisker plots illustrating the fraction of time animals spent in three different states: motion, rest and transition. The plot shows representative data from one mouse (same as in ***A***, n=3 consecutive 10-min-long image series). ***D***, Same analyses as in ***C*** but the plot shows a median of 7 mice. A significant difference was observed between the fractions of time spent in 3 different conditions (One-way repeated measure ANOVA followed by Holm-Sidak multiple comparisons test, 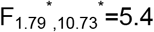, P = 0.03, here and below a star (^*^) denotes Greenhaus-Geisser sphericity correction). However, pairwise comparisons using Holm-Sidak post-hoc test did not reach the level of statistical significance. ***E***, Box-and-whisker plot illustrating spatial correlation coefficients within and between maps obtained during motion and resting time periods (3 maps, each representing a 10-min-long image series recorded in a mouse shown in ***A***). ***F***, Box-and-whisker plot illustrating median (per mouse) spatial correlation coefficients within and between maps obtained during motion and resting time periods. Obtained values are significantly different (One-way repeated measure ANOVA followed by Holm-Sidak multiple comparisons test, 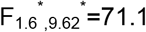, P < 10^−3^, motion vs. rest: P = 10^−3^, the other two comparisons: P < 10^−3^).

In neonatal rodents, we and others have characterized multiple types of intrinsic cortical activity patterns with distinct spatiotemporal properties. *In vivo,* the latter include spindle bursts in the developing visual and somatosensory cortex (Hanganu et al., 2006; Khazipov et al., 2004), gamma and spindle bursts in the primary motor cortex (An et al., 2014), spindle-shaped field oscillations in the prefrontal cortex (Brockmann et al., 2011), intermittent population bursts in the medial entorhinal cortex (Valeeva et al., 2019), early sharp waves in the hippocampus (Valeeva et al., 2019), cortical Ca^2+^ waves/transients and accompanying spindle-shaped oscillations in the temporal and occipital cortex (Adelsberger et al., 2005; Kirmse et al., 2015), etc. Similar patterns of early spontaneous cortical activity were also found in preterm infants (McVea et al., 2016; Vanhatalo et al., 2002). In the latter, abnormal activity patterns were associated with adverse cognitive outcomes (Arichi et al., 2017), whereas the frequency of physiologically patterned activity was positively correlated with brain growth (Benders et al., 2015).

So far, however, the mentioned above activity patterns were mostly studied in isolation, from the perspective of a given sensory or sensorimotor system (Ackman and Crair, 2014; An et al., 2014; Anton-Bolanos et al., 2019; Khazipov and Milh, 2017). The results of these studies suggest that spontaneous activity of the peripheral sensory organs (e.g. developing retina or cochlea) or spontaneous muscle twitches interact with intrinsic cortical oscillators thereby modulating neuronal spiking in corresponding cortical regions (Luhmann et al., 2016). Moreover, the spontaneous network activity in some cortical regions (e.g. prefrontal, motor, or somatosensory cortex) was reported to be entrained by the activity in the other cortical/subcortical structures (e.g. hippocampus, S1, thalamus, or M1, respectively) likely priming the functional coupling between the two areas seen in the adulthood (An et al., 2014; Anton-Bolanos et al., 2019; Brockmann et al., 2011). Similarly, spontaneous network activity was suggested to sculpt both inter- and intra-hemispheric efferent projections from the developing visual cortex of rats and ferrets (Ackman and Crair, 2014). Together, this data suggests that early cortical development is driven by highly complex and hierarchic patterns of endogenous activity. However, the integral map of intrinsic cortical activity in neonates as well as the spatiotemporal relationship between the activity patterns in different cortical areas remain obscure.

To clarify these issues, we captured intrinsic cortical activity by means of large-scale single-photon Ca^2+^ imaging and analyzed it using data-driven mathematical techniques. The data-driven nature of our approach precluded the use of the standard region of interest (ROI)-based spatial segmentation technique, as the spatial location of the endogenous activity patterns was a priory unknown. Forcing the activity pattern to be confined to a given anatomical region could be meaningful for sensory-driven but not for spontaneous activity. Moreover, this approach does not apply to the developing brain where there is no fine-grained anatomical atlas and activity patterns change on a day-to-day basis, due to maturation of the underlying neural circuitry. Thus, automatic feature selection techniques, able to extract relevant features of the original high dimensional data space, had to be used. We chose a decomposition technique (Non-negative Matrix Factorization (NMF)), which doesn’t need prior information about the location of ROIs and extracts information about activity patterns by positively constraining spatial and temporal components and relaxing the orthogonality constrain (used e.g. in Principal Component Analysis (PCA)), which is not essential. To study functional connectivity (i.e. statistical measure of synchronicity between activities in two different ROIs), we calculated the inverse covariance matrix and penalized our model with a sparsity regularizer (Friedman et al., 2008).

By using the described above mathematical workflow (schematically illustrated in Fig. 2-1), we characterized the self-initiated motion- and rest-specific spatiotemporal activity patterns in the dorsal cortex of neonates. The developmental stage (P3), analyzed in this study, is reminiscent of late gestation (preterm) stage in human babies, when (i) long-range neuronal projections are still developing, and (ii) the electrical brain activity consists of developmentally unique, intermittent events, believed to guide activity-dependent brain wiring (Omidvarnia et al., 2014; Vanhatalo et al., 2002).

## Materials and methods

### Mice

All experiments were conducted in accordance with institutional animal welfare guidelines and were approved by the state government of Baden-Württemberg, Germany. Because during the first postnatal week the rapidly developing rodent cortex is known to traverse several developmental states (Kirischuk et al., 2017), for the sake of consistency we focused on a defined animal age (P3), characterized by cortical early network oscillations (Adelsberger et al., 2005; Garaschuk et al., 2000; Kirmse et al., 2015) with their developmentally unique intermittent activity, first appearance of spontaneous activity patterns in the prefrontal cortex (Brockmann et al., 2011) but immature callosal projections (Son et al., 2017; Wang et al., 2007). In accord with mentioned above human studies (Omidvarnia et al., 2014), we use the term network as a functional concept referring to a set of brain regions displaying simultaneous activity or a high degree of functional connectivity, as measured by the mathematical paradigms used.

In line with the biometric planning, seven 3-day-old nestin-Cre × Ai95(RCL‐GCaMP6f)-D and three age-matched wild type C57BL/6) mouse pups of either sex were used. Parent mouse lines B6.Cg-Tg(Nes-cre)1Kln/J and B6;129S-Gt(ROSA)26Sortm95.1(CAG-GCaMP6f) Hze/J were originally obtained from Jackson Laboratory (stock № 003771 and stock № 024105, respectively) and were bred on the C57BL/6 background.

### Animal preparation for in vivo Ca^2+^ imaging

Mouse pups were anesthetized with isoflurane (2.5% for induction, 1-2% for the rest of surgery). The skin above the dorsal part of the skull was cut away, the connective tissue was gently peeled off and the custom-made ring-like plastic chamber was glued to the skull. Xylocaine gel (2%) was applied to the wounded skin edges. To prevent the skull from drying out, it was covered with agarose (1-2%) dissolved in standard Ringer’s solution containing (in mM): NaCl 125, KCl 4.5, MgCl_2_ 1, CaCl_2_ 2, NaHCO_3_ 26, NaH_2_PO_4_ 1.25, Glucose 20, pH 7.4. The procedure described above lasted 23 ± 3.8 min (n=7 mice). Thereafter, the isoflurane anesthesia was terminated and the animal was allowed to recover on a warming plate (34°C - 36°C) for at least one hour. This well established (Ackman et al., 2012; Adelsberger et al., 2005; Che et al., 2018; Dooley et al., 2020; Hagihara et al., 2015) protocol provided stable recording conditions with similar ongoing *in vivo* activity patterns, observed within and across experimental animals (Fig. 2-2).

### In vivo large-scale single-photon Ca^2+^ imaging

After recovery the animal, resting on a warming plate, was transferred into the imaging setup and head-fixed under the MVX10 Research Macro Zoom Microscope equipped with an LED source for excitation (Thorlabs, central wavelength 470 nm), 470/40 nm excitation filter, 495LP beam splitter, 525/50 nm emission filter (all from Chroma Technology), the Zyla 4.2 sCMOS camera (Andor Technology) and Andor Solis software for image acquisition. Animal’s limbs were free to interact with each other and with the surface of the warming plate. Both the left and the right hemispheres were imaged simultaneously through the intact skull at 256×256 pixel resolution (1 pixel ≈ 25×25 μm^2^) and 11 ms/frame for half an hour (three consecutive 10-min-long acquisition series). During the recordings, mice spent 70.6 ± 12% of their time in a state without any apparent movement (Fig. 2C and D). This behavior is very similar to that of control neonatal mice (Fig. 3e in (Adelsberger et al., 2005)) and differs substantially from the one indicating distress (e.g. sudden movements, vocalization, and urination) (Castelhano-Carlos et al., 2010).

**Fig. 3.**
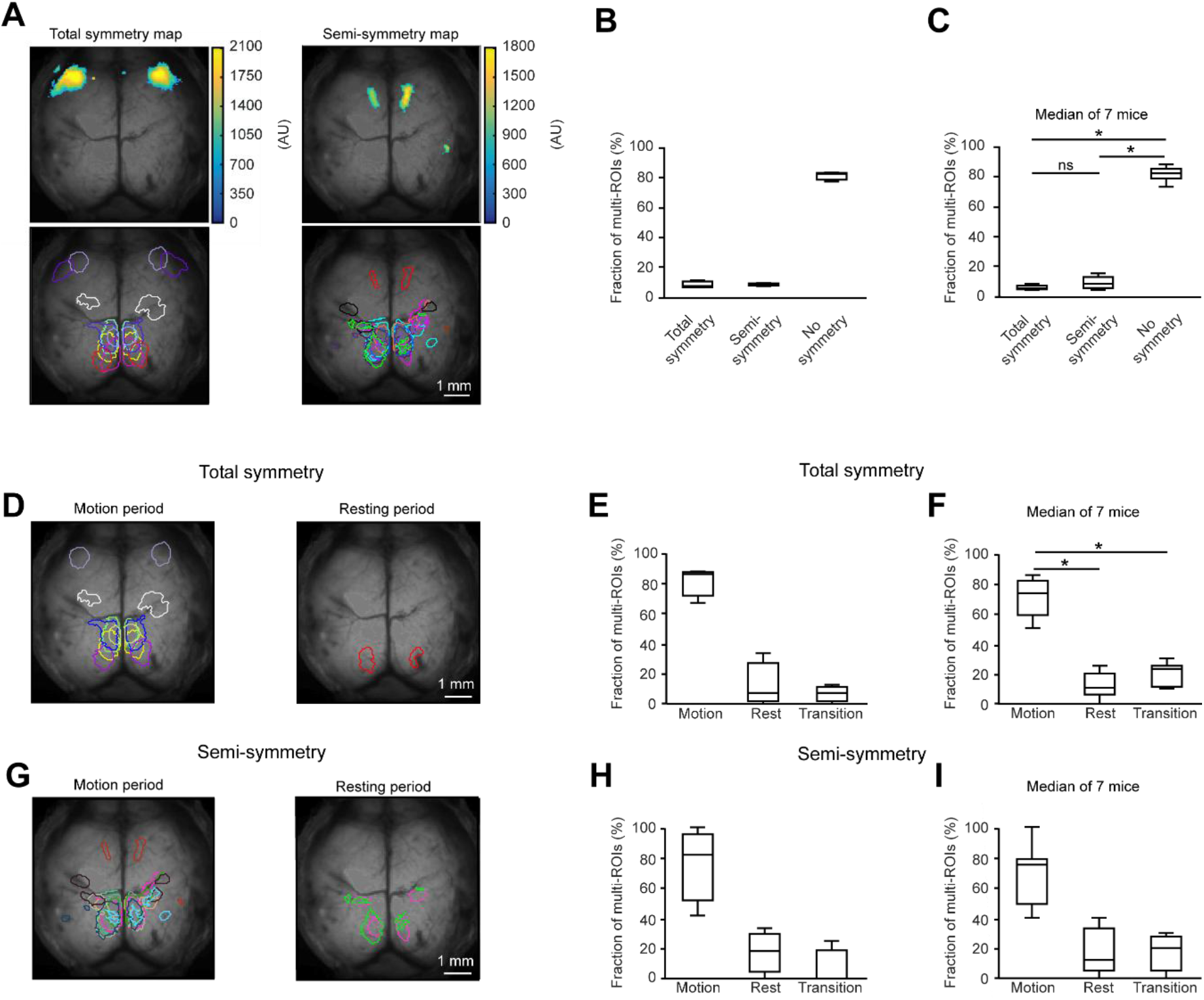
Hemisphere-symmetric neuronal activity in the neonatal mouse cortex. ***A***, Representative active subregions belonging to one multi-ROI filter (upper) as well as symmetry maps (lower) projected on a grayscale image of P3 mouse cortex. Here and in Fig. 4 the sample data are from mouse 4 (Fig. 2-2). Upper panels: fluorescence signals are color-coded with warmer colors indicating higher signal intensity. Lower panels: different colors delineate multi-ROIs, each satisfying the total (left panel) or semi- (right panel) symmetry criterion (see Materials and methods). ***B***, Box-and-whisker plot illustrating the median fractions of multi-ROIs in three distinct categories: total, semi-, and no symmetry (3 consecutive 10-min-long image series recorded in mouse 4). ***C***, Same analyses as in ***B*** illustrating the median data obtained in 7 different animals. Obtained values are significantly different (One-way repeated measure ANOVA followed by Holm-Sidak multiple comparisons test, 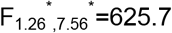, P < 10^−3^; P < 10^−3^ for both semi-symmetry vs. no symmetry, and total symmetry vs. no symmetry comparisons). ***D***, Representative total symmetry maps recorded during motion (left panel) and resting (right panel) time periods (mouse 4). Different colors delineate multi-ROIs, each satisfying the total symmetry criterion. ***E***, Box-and-whisker plot showing the median fractions of symmetric multi-ROIs during the three time periods: motion, rest, and transition (3 consecutive 10-min-long image series recorded in mouse 4). ***F***, Same analyses as in ***E*** illustrating the median data obtained in 7 different animals. Obtained values are significantly different (One-way repeated measure ANOVA followed by Holm-Sidak multiple comparisons test, 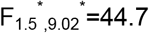, P < 10^−3^; P = 10^−3^ for both comparisons). ***G***, Similar maps as in ***D*** but here colors delineate multi-ROIs, each satisfying the semi-symmetry criterion. ***H***, Box-and-whisker plot showing the median fractions semi-symmetric multi-ROIs during the three time periods: motion, rest, and transition (3 consecutive 10-min-long image series recorded in mouse 4). ***I***, Same analyses as in ***H*** illustrating the median data obtained in 7 different animals.

#### The nature of recorded Ca^2+^ signals

To understand the nature of the recorded large-scale Ca^2+^ signals we imaged them in the temporal cortex using high resolution two-photon microscopy in brain slices because this cortical region shows similar spontaneous large-scale Ca^2+^ signals *in situ* and *in vivo* (Adelsberger et al., 2005). To do so, at the end of *in vivo* imaging experiments horizontal 500-μm-thick slices were prepared from three P3 nestin-Cre × Ai95(RCL‐GCaMP6f)-D mice as described earlier (Adelsberger et al., 2005). As the nestin promotor, driving the Cre recombinase activity in our mice, is expressed in both neuronal and glial precursors, both neurons and astrocytes expressed GCaMP6f (Fig. 2-3). In addition, astrocytes were stained by bathing the slice for 10 min at 37 °C in the standard Ringer’s solution (composition see above) containing the red fluorescent dye sulforhodamine 101 (S101, 3 μg/ml; Fig. 2-3A) (Adelsberger et al., 2005). To speed up the data acquisition, we selected fields of view with a rather high density of astrocytes. Ca^2+^ imaging was performed at 32-34 °C using a two-photon microscope (Olympus Fluoview 1000, Olympus) coupled to a mode-locked Ti-Sapphire laser working at a wavelength of 710-990 nm (MaiTai, Spectra Physics) and equipped with a water-immersion objective (Olympus 60x, 1.0 NA). The emission signals of GCaMP6f and S101 were split by a 570 nm dichroic mirror and sent through the BP 536/40 and BP 630/92 filters, respectively. We analyzed ROIs containing individual cells (neurons and astrocytes) as well as the large-scale ROIs (176 × 176 μm), covering the entire imaging frame. The latter contained the mixed signal alike our *in vivo* data. ROIs for each individual neuron or astrocyte were drawn manually. The respective traces were calculated by averaging all pixels inside the ROI. Ca^2+^ signals were detected by the Matlab (R2016b) peak detection function after smoothing original traces (moving average function, 1 s moving window size). As shown in Fig. 2-3B, the Ca^2+^ signals recorded from large-scale ROIs closely resembled signals of individual neurons, while astrocytes either showed no Ca^2+^ signals at all (88 of 119 analyzed cells) or showed Ca^2+^ signals with much slower kinetics (31 of 119 analyzed cells; Fig. 2-3C). Note that in astrocytes the frequency of spontaneous Ca^2+^ signals is much lower than in neurons (Adelsberger et al., 2005; Nimmerjahn et al., 2004; Tian et al., 2006), thus explaining the rare appearance of the latter in our recordings. The similarity among traces was determined using the Pearson correlation coefficient. Correlation coefficients among neurons and between each neuron and the large-scale ROI were high, quite opposite to those obtained when correlating the respective astrocytic data (Fig. 2-3D). The median correlation coefficients obtained when comparing individual neurons with the respective large-scale ROIs were as high as 0.9 ± 0.1 (n=190 neuron/large ROI pairs; Fig. 2-3E). The much lower coefficients were obtained when conducting the same comparison for astrocytes (0.29 ± 0.27, n=119 astrocyte/large ROI pairs).

We concluded, therefore, that *in vivo* Ca^2+^ signals analyzed in the present study are of neuronal origin. This conclusion is further supported by the fact that between P0 and P3 neurons represent ~ 88% of all cells in the rodent cortex (Bandeira et al., 2009).

### Recordings of animal’s movement

Animals were imaged with a monochrome infrared (IR) light-sensitive camera. IR LED (950 nm) was used for illumination. To remove the salt-and-pepper noise, the image sequence (720×480 pixels, 29 Hz) was filtered with a median filter of kernel size 3 × 3. Subsequently, adjacent frames were pixel-wise subtracted from each other thus transforming the original image sequence into an image sequence containing the information about the temporal changes within the field of view (Lipton et al., 1998). For a given resulting image, the sum of absolute values of all pixels was taken as a measure of instantaneous body motion. Subsequently, the movement trace was upsampled using Matlab’s 1D interpolation function to match the sampling rate of the Ca^2+^ signals (~91 Hz), plotted over time, and normalized to the maximum recorded value. The breathing-related motion artifacts were identified by the back-to-back comparison of the animal behavior and the corresponding movement trace. For each recording, the threshold was selected such that these artifacts were discarded. On average the threshold was ~6% of the maximum recorded value and was used to create a binary signal, where all ones corresponded to the motion period and all zeros corresponded to the resting state (Fig. 2-4). The on/off switching of the 470 nm excitation light, recorded by both cameras, was used to synchronize the imaging of Ca^2+^ and movement signals. In addition, we defined the time window including one second before the movement onset and three seconds after the end of the movement (corresponding to the decay time of the eventual movement-evoked Ca^2+^ signals) as a transition state. With this procedure, the behavior of the animal was automatically subdivided into three different states: motion, rest, and transition.

As the motion state potentially contained either spontaneous muscle twitches (Blumberg, 2010) or generalized movements, in a separate series of experiments we recorded nuchal muscle electromyogram (EMG (Seelke et al., 2005)) simultaneously with imaging body movements. Note that at P3 the EEG traces are discontinuous thus providing no information about the vigilance states (Blumberg et al., 2014; Jouvet-Mounier et al., 1970; Rensing et al., 2018). For EMG recordings a stainless steel wire (type SS-3T, Science Products GmbH, Hofheim, Germany) was used. Two to five mm of the insulation were stripped off and the insulation-free ends of the wire were gently inserted subcutaneously and placed bilaterally on the top of the neck muscles. After the correct position of the wires has been verified, the wires were fixed to the ring-like plastic chamber (see above) with UV-cured dental cement. EMG data were recorded at 1 kHz (0.3-500 Hz band-pass filter) using the Powerlab differential amplifier (ADInstruments Ltd, Oxford, United Kingdom).

Movement binary signal was created from the EMG signal using signal envelope analysis (Oppenheim et al., 1999). The normalized cross-correlation between the movement signals extracted from the imaging and the EMG data showed peak values of 0.6-0.85 (n = 9 recording in 3 mice) suggesting that both signals reflect the structure of the animal’s movement in a consistent way. Note, however, that some tiny movements, as well as limb twitches, likely remain unrecognized by either one or even both techniques (Seelke et al., 2005). This fact and the fragmentation of sleep in the neonatal mice (Fig. 2-4, see also Fig. 3A in (Blumberg et al., 2014)) made unequivocal identification of active and quiet sleep periods difficult. Next, we separately used EMG- and imaging-based data sets of individual animals to construct the distributions of the durations of movement episodes. Both data sets identified the first peak at 350-600 ms, followed by a local minimum at 700-800 ms. Based on this data we chose 750 ms as an empirical border between spontaneous muscle twitches and generalized movements. In our hands, muscle twitches represented 39.5 ± 14 % of all movement events (n=6 mice) and the fraction of time covered by muscle twitches ranged between 5% and 16% (median per mouse) of the total movement time. We concluded, therefore, that under our experimental conditions Ca^2+^ signals recorded during the animal’s movement mainly reflect the ones associated with generalized movements.

### Analyses of spontaneous cortical activity

If not otherwise indicated, data analyses started with reducing the image size to 128×128 pixels using 2 × 2 binning and motion correction (Matlab R2016b image registration toolbox; Fig. 2-1). We used full-band data for all analyses, no frequency filtering was applied. Because our (see Movie 2-1 recorded in WT mice) as well as literature (Kozberg et al., 2016) data show that localized neural activity in neonatal mice does not evoke local functional hyperemia (in contrast to the adult brain), no hemodynamic correction was applied.

#### ROI detection based on Nonnegative Matrix Factorization (NMF)

Pixels that showed a coherent change in fluorescence intensity were grouped into ROIs using the NMF algorithm, running on a full data set. According to NMF, a given data matrix X is factorized into two matrices *W* and *H* containing spatial filters (*W*) and the corresponding time information (*H*): X = *WH*, *W*_*ik*_ > 0 & *H*_*kj*_ > 0. *W* and *H* can be calculated by minimizing a cost function, *min*_*W,H*_∥*X* − *WH*∥_F_. In the case of the low-rank NMF, the rank (k) of *W* and *H* is set to be smaller than the dimension of X. To minimize the cost function we have chosen the alternating least-squares algorithm (Berry et al., 2007). The initialization method was based on ICA (Fig. 2-1). Selection of the rank for *W* and *H* relied on PCA singular values. We found that selecting 300 components in the PCA insured that the explained variance in the data was ≥95 % whilst data noise was significantly reduced.

The 300 spatial filters were inspected by eye. Filters resembling either the blood vessel pattern or the whole brain image contaminated by blood vessels (e.g., Figs. 2-5A and 2-5B, Movies 2-1 and 2-2), likely reflecting movement artifacts, were discarded. The total number of such filters was in the range of 95 ± 34. The total data variance, explained by the discarded data, amounted to 36 ± 3% (mean ± S.E.M, n=7 mice). The remaining filters contained groups of active neighboring pixels, clearly discernible from the noisy background (e.g., Fig. 2-5C). Sobel edge detector algorithm (Lim, 1990) was used to define the border between the active and background pixels. Finally, the filters were binarized using the threshold value of fourfold the SD of the corresponding background noise (see above). Groups of connected pixels with n > 10 pixels were defined as ROIs. The filters containing one ROI were called **single ROI filters** and the ones with more than one ROI were called **multi-ROI filters**. To create an ROI-based frequency map, each ROI was weighted by ascribing a value equal to the number of peaks detected in its temporal domain (*H*). For peak detection traces corresponding to spatial filters (*W*) were thresholded at 95% of their maximal amplitude. Finally, the weighted ROIs were superimposed to create a color-coded ROI-based frequency map (e.g. Fig. 2A).

#### Calculating the association between fluorescence intensity in C57BL/6 mice and the body movement

Using the result of NMF analysis, the data was reconstructed based on those spatial filters which only contain the blood vessel pattern or the whole brain image contaminated by the blood vessel pattern (exemplified in Figs. 2-5A and 2-5B), likely reflecting movement artifacts. Using the reconstructed video, the trace of the global fluorescence intensity was calculated by averaging the intensity of all pixels within the recorded brain area. Taking this trace, the Gaussian mixture model with 2 Gaussians was used to separate large intensity changes from small baseline fluctuations. The intercept of two Gaussians was taken as the separation threshold. Next, the calculated trace was binarized using the threshold value, returning a sequence of individual blocks of activity. Finally, the association between the binary fluorescence intensity signal and the binary movement signal was calculated using the following formula:

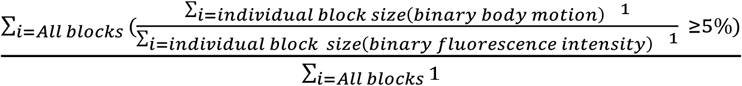

#### Symmetry maps

For calculation of the cortical symmetry maps, one hemisphere was mirrored to another against the mid-sagittal plane and all ROIs belonging to each multi-ROI filter were examined pair-wise by calculating their two-dimensional correlation coefficients. A pair of ROIs was considered symmetric if the correlation coefficient was ≥0.5. Spatial filters which contained only symmetric ROIs were classified as **total symmetry filters** and filters with at least one pair of symmetric ROIs were classified as **semi-symmetry filters**. Further, according to their corresponding time information (matrix *H*) and the animal state, the symmetry filters were subdivided into three categories: motion, rest, and transition. Superpositions of filters of a given class and category were used to produce behavioral state-specific symmetry maps.

#### Wave analyses

To detect propagating Ca^2+^ waves the image sequences were processed by the nonparametric spectral estimation method: Multi-channel Singular Spectrum Analysis (MSSA; (Golyandina and Usevich, 2010)). Before being reshaped into a two-dimensional space, full-band data were down-sampled by a factor of 10 in the temporal domain. The resulting matrix *D*_*n×p*_ was centered in the time domain by subtracting the mean value of each pixel. Thus,

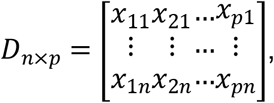

where *n* is the number of samples in time and *p* is the number of pixels in each image.

According to the MSSA algorithm, we determined the embedding matrix M and covariance matrix *C* as 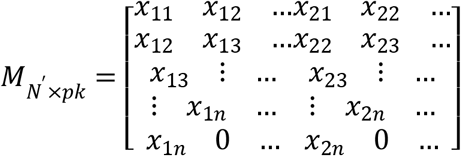, where *N*^′^ = *n* − *k* + 1, and *k* is a lag shift.

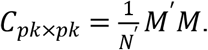

By eigen decomposition of the covariance matrix *C*, principle vectors and principle components were calculated. In the MSSA, each principle vector describes an oscillatory component in the data set whereas principle components represent the corresponding weighting coefficients. The k value (lag shift) was set to 15 s.

Data were reconstructed (MSSA (Golyandina and Usevich, 2010)) based on a subgroup of components (sorted components 20-150^th^). The remaining components represented either noise or slow baseline drifts. Finally, the reconstructed data were spatially thresholded to obtain the corresponding binary mask of each frame. Thresholding was based on fitting Laplace distribution to the reconstructed data. As a threshold, we selected a fourfold of scale parameter of the estimated Laplace distribution. The location, in which the activity was observed in the first frame of each individual wave, was defined as its pacemaker region. Populations of connected pixels were treated as objects and dynamics of each object in time represented a single wave. The distance traveled by each wave was characterized by the sum of displacements of the corresponding object’s center of mass. The waves were divided into two groups: (1) stationary waves with the center of mass moving less than 200 μm and (2) propagating waves with the center of mass moving more than 200 μm. To calculate the speed of a wave we divided the total distance traveled by the respective time. Further, we counted the waves propagating within a given cortical region and calculated the fraction of waves occurring during the motion and resting periods, respectively. A similar procedure was applied to calculate the fraction of waves propagating between any two cortical regions. One animal (Mouse 6 in Fig. 2-2) was excluded from the above analyses because compared to other mice, this data set contained more brief motion periods disrupting the resting period and thus making the analysis of wave propagation and functional connectivity impossible.

### Map of simultaneously active cortical regions

To identify brain regions synchronously active during the movement and the resting states, spatial filters in multi-ROIs obtained from NMF analysis (see above) were processed according to the time information (matrix *H*) and the animal state. Then, the subregions from each multi-ROI were assigned to the anatomical regions based on the location of their center of mass. For any multi-ROI filter, each pair of simultaneously active subregions (within each cortical region or between the two distinct cortical regions) was represented by an entry in the matrix (10 × 10 - each row or column corresponds to one of the cortical regions (see Fig. 5-1)). Such entry was made each time when the two subregions were simultaneously active. This procedure was repeated for all multi-ROIs in motion and resting states providing two matrices. Then each matrix was normalized to the sum of both matrices and values below 50% were removed, to focus on interactions, predominant for each state. The median of three matrices (three 10-min-long image series) was taken as a representative map for each mouse (Fig. 5-1C) and the median of all maps from n = 6 mice was depicted in Fig. 5-1D.

**Fig. 5.**
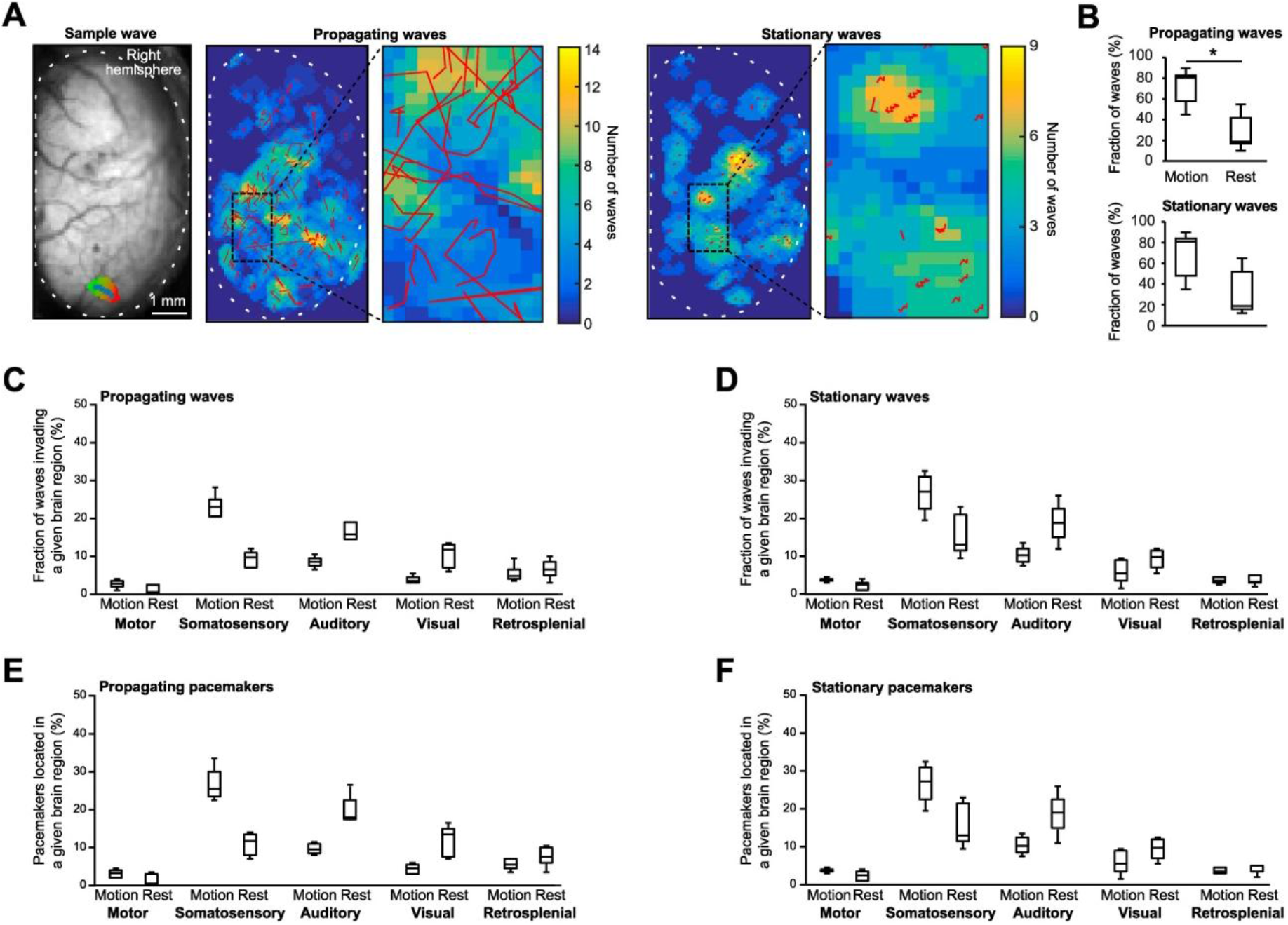
Stationary and propagating Ca^2+^ waves in the neonatal mouse cortex. ***A***, Left panel shows (from left to right): (i) top view on a right hemisphere of P3 mouse cortex (mouse 5 in Fig. 2-2) with a sample propagating wave (the direction of movement is color coded from green to red, blue stars depict the location of the center of mass in each frame (see Materials and methods); (ii) a superposition of propagating waves recorded in this animal in 6 min (here and below the corresponding center of mass trajectories are depicted as red lines) and (iii) the framed area, shown on an expanded scale. Right panel shows (from left to right): (i) a superposition of stationary waves recorded in this animal in 6 min and (ii) the framed area shown on an expanded scale. Broken lines delineate the contour of the brain area imaged in this experiment. False colors reflect the number of waves each pixel was involved into. ***B***, Box-and-whisker plots showing median (per mouse) fractions of propagating (upper panel - t_5_ = 3.21, P = 0.024) and stationary (lower panel) waves happening during motion and rest, respectively (n = 6 mice). For each mouse, the number of propagating waves during the motion (or rest) was normalized to the number of all propagating waves recorded. A similar normalization procedure was used for stationary waves. ***C***, ***D**,* Box-and-whisker plots showing averaged between the two hemispheres median (per mouse) fractions of propagating ***C***, and stationary ***D***, waves invading the respective cortical regions. ***E***, ***F**,* Similar analyses as in ***C**, **D*** but illustrating median (per mouse) fractions of pacemakers for propagating ***E***, and stationary ***F***, waves located in the given cortical region (n = 6 mice). To calculate the fraction of propagating/stationary waves in a given cortical region during the motion/rest the number of propagating waves observed in this region during this particular state was normalized to the number of all propagating waves detected in this state.

### Functional connectivity maps

For calculating direct connectivity maps, input data were binned 5 by 5 pixels, temporally down-sampled by a factor of 10, and centered in the time domain by subtracting the mean value of each pixel. Thereafter, the connectivity maps were calculated separately for resting and motion periods by using the sparse partial correlation algorithm (Smith, 2012; Smith et al., 2011). Assuming that data are drawn from a multivariate Gaussian distribution, elements of the precision matrix Σ^−1^ explain partial covariance in a given data set. Sparsity constraint was introduced by adding an extra penalty term to the likelihood function. To calculate the sparse partial covariance one has to maximize the penalized log-likelihood L (Friedman et al., 2008),

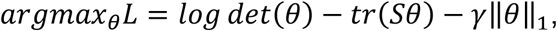

where θ = Σ^−1^, and *S* is the empirical covariance matrix ∥ ∥_1_ is |_1_ norm and γ is a sparseness tuning parameter.

The sparse partial covariance was calculated with a range of different γ values. The smaller γ values provide a more dense connectivity pattern, whereas the larger γ values increase sparseness in the connectivity map by removing weak connections between the nodes. After obtaining the direct connectivity map, pixels belonging to specific cortical regions were identified using the corresponding anatomical map of the brain (like the one shown in Fig. 1A). Partial covariances between pixels of each specific cortical region were summed up giving the value α_ij_, where α_ij_ reflects the strength of the connectivity within the given cortical region if i=j. Similarly, partial covariances between pixels belonging to a pair of specific cortical regions were summed up, contributing those values of α_ij_ (i≠j), where α_ij_ reflects the strength of the connectivity between the two different cortical regions. Finally, all values were normalized to the maximal α_ij_ value.

**Fig. 1.**
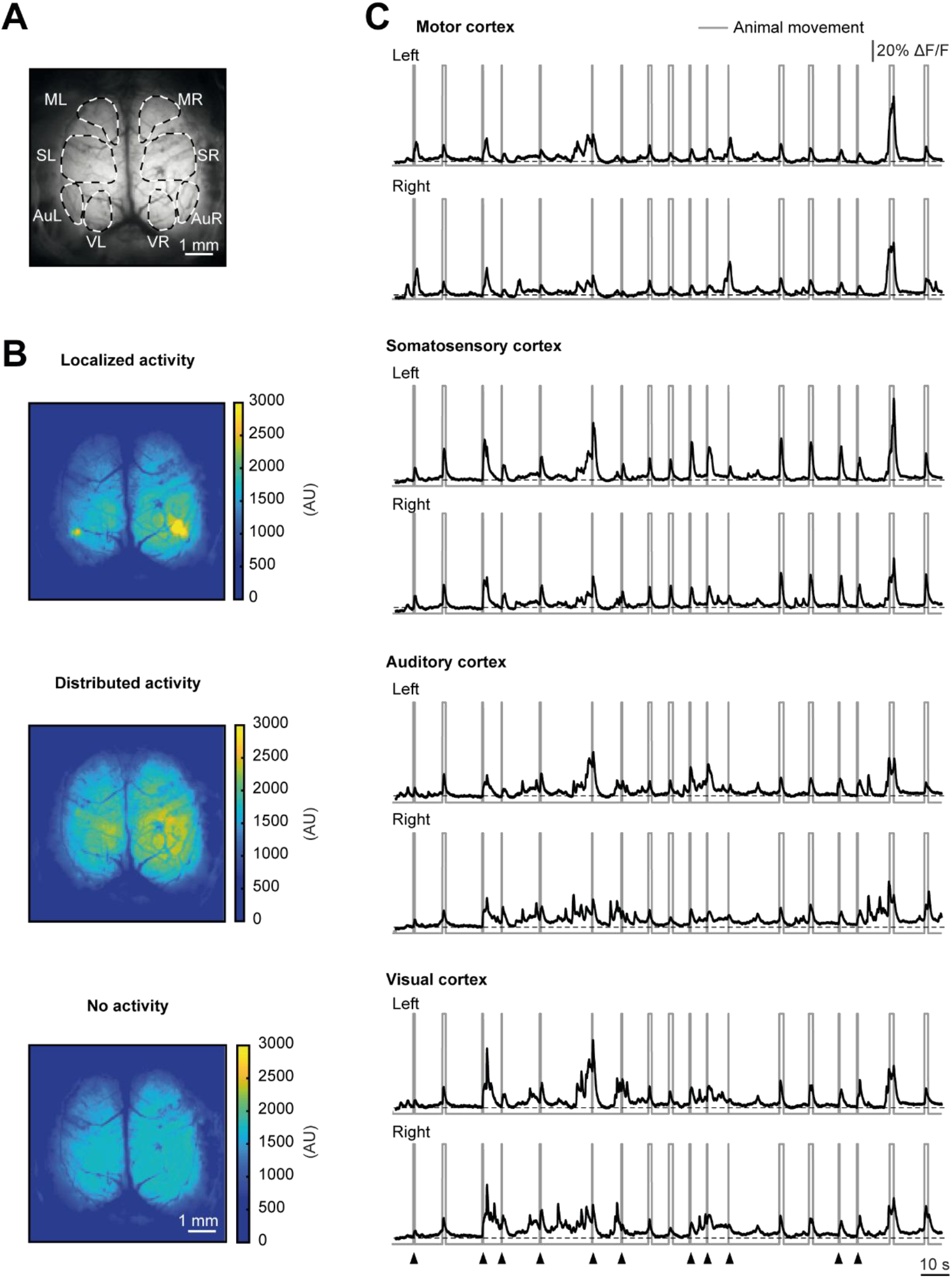
Large-scale spontaneous neuronal activity in the neonatal mouse cortex. ***A***, Top view on a P3 mouse cortex taken through an intact skull. Broken lines delineate cortical regions of interest (estimated as in (Gee et al., 2014)): ML, MR - motor cortex (left and right), SL, SR - somatosensory cortex (left and right), AuL, AuR - auditory cortex (left and right) and VL, VR - visual cortex (left and right). ***B***, Averages of 100 consecutive autofluorescence-subtracted images taken during periods of either local (top) or global (middle) cortical activities or no activity (bottom). Autofluorescence values are averages of mean pixel values recorded in three different 3-day-old C57BL/6 mice. Subtraction was done for display purposes only. Image brightness (arbitrary units, AU) is color coded, with warm colors reflecting higher values. ***C***, 200-second-long ΔF/F traces recorded from ROIs delineated in ***A***. Gray boxes mark periods of animal’s movement. Arrowheads mark muscle twitches. Data shown are from mouse 1 (Fig. 2-2).

### Statistical analyses

Statistics was performed using JASP and SPSS software. The normality of the data distribution was tested using the Shapiro-Wilk test. The two-tailed paired Student’s t-test was used for pair-wise comparisons of normally distributed data sets. For pair-wise comparisons of not normally distributed data, Wilcoxon signed-rank test was applied. One-way or two-way repeated measures ANOVA (rANOVA) was used for comparing more than two normally distributed dependent variables. In the case of not normally distributed data, the Friedman test was used. In rANOVA sphericity assumption was tested using Mauchly’s test and in case of sphericity violation, Greenhouse-Geisser correction was applied. All ANOVAs, which show significant results, were followed by a post-hoc test with Hold-Sidak correction for multiple comparisons. For all tests, differences were considered significant if P < 0.05. If not otherwise indicated, data are presented as median ± interquartile range (IQR)).

## Results

To monitor patterned neuronal activity in the dorsal cortex of neonates, we imaged non-anesthetized nestin-Cre × Ai95(RCL‐GCaMP6f)-D mice through an intact skull with a high speed/ high quantum efficiency sCMOS camera. Although the use of genetically-modified mice is the only technique enabling large-scale cortical Ca^2+^ imaging in this young age, the use of Cre-driver lines was recently questioned by the work of Steinmetz et al., reporting the presence of the epileptiform activity in different crosses of Ai93/Ai94 and various Cre-driver mice (Steinmetz et al., 2017). Unlike the Ai93/Ai94 mice, Ai95 mice used here do not contain tetracycline-controlled transactivator protein (tTA), reported to be neurotoxic (Han et al., 2012). Still, we searched for non-physiological activity in our mice using the same approach as Steinmetz et al. We analyzed fluorescence traces recorded in the frontal and the visual cortices and only found events with variable amplitude, duration, and shape (Fig. 1-1), in sharp contrast to large amplitude, brief duration, stereotyped shape epileptiform events recorded by Steinmetz et al. (see their Figs. 2, 3). The fact that we did not observe any pathological activity is consistent with the literature, showing that in used here B6.Cg-Tg(Nes-cre)1Kln/J mice expression of the Cre recombinase becomes widespread only during the perinatal development (around P0; (Liang et al., 2012)). With data acquired at P3, this leaves any pathology no time to develop. Moreover, Steinmetz et al. never saw any aberrant activity before the age of at least 7 weeks, again arguing against the presence of any pathology at P3.

### Patterns of spontaneous activity in the dorsal cortex of neonates

As shown in Fig. 1A-C, we observed diverse patterns of endogenous activity ranging from localized signals, involving one or several anatomical regions (Fig. 1B, upper panel) to large distributed signals, covering almost the entire dorsal cortical surface (Fig. 1B, middle panel). Placing regions of interest onto major anatomical regions of the dorsal cortex (i.e. visual, auditory, somatosensory, and motor cortices) revealed rich patterns of recurring changes in ΔF/F in each studied region (Fig. 1C). Simultaneous recording of animal movements (consisting at this age of muscle twitches and generalized movements, see Materials and methods for details about the estimation of movement’s nature) by means of an infrared imaging camera (a gray trace in Fig. 1C; Fig. 2-4), revealed the presence of changes in ΔF/F during both moving and resting states.

To understand the nature of the observed changes in ΔF/F, pixels that showed a coherent change in fluorescence intensity were grouped into ROIs using NMF (see Materials and methods), running on a full 10-min-long set of data. The analyses returned 3 typical ROI patterns (so-called spatial filters) resembling (i) blood vessel pattern (Fig. 2-5A), (ii) the whole brain image contaminated by the blood vessel pattern (Fig. 2-5B), or (iii) groups of active neighboring pixels, clearly discernible from the background (Fig. 2-5C).

The first two ROI patterns were also seen in wild type C57BL/6 mice (n=3), where 95% of them were associated with animal’s movement, whereas the third one was observed only in GCaMP6f-expressing nestin-Cre × Ai95(RCL‐GCaMP6f)-D mice (compare Movies 2-1 and 2-2). Thus, although distributed fluorescence signals, seen in nestin-Cre × Ai95(RCL‐GCaMP6f)-D mice (Fig. 1B, middle panel), likely also include a Ca^2+^-sensitive component, in-depth analysis of these events was precluded by contaminating movement artifacts and, if any, artifacts, related to flavoprotein autofluorescence and confounding hemoglobin absorption.

### Behavioral state-dependent patterns of local cortical activity

In the subsequent analyses, we thus concentrated on the properties of localized activity, which mainly reflected Ca^2+^ signaling in the underlying neuronal population (Fig. 2-3, see Materials and methods).

Regarding the localized activity, the NMF algorithm identified either “single ROIs”, in which all coherently active pixels were immediately adjacent to each other, or “multi-ROIs”, in which coherently active pixels were distributed in patches throughout the dorsal cortex (see Materials and methods for details). Next, we constructed ROI-based frequency maps of the local spontaneous activity during the motion and the resting periods (Fig. 2A and B; see Materials and methods for details). To clearly discriminate between the motion and the resting states we separated them by transition periods including one second before the movement onset and three seconds after the end of the movement (Fig. 2C and D). Analysis of the amount of variance (Pedregosa et al., 2011) in the data related to each state (rest, transition (before and after the movement), generalized movements, and muscle twitches) showed that the amount of variance related to twitches was 0.2 ± 0.6% and thus much less than that related to generalized movements (61.98 ± 9.39 %; Fig. 2-6), probably due to the short duration of twitches (see Materials and methods). Therefore, the motion-related frequency maps almost exclusively explain the activity, happening during the generalized movements. Overall, the 3-day-old mice moved only 32 ± 18.5% of the recording time (Fig. 2D), thus spending most of their time in a state with no detectable motion. This is consistent with our earlier data described in ref. (Adelsberger et al., 2005).

On average, coherently active local cortical subregions belonging to either single or multi-ROIs covered the areas of approximately 0.2 mm^2^ (Fig. 2-7) and the size of these areas was similar during different behaviors (rest: 0.21 ± 0.06, generalized movements: 0.17 ± 0.03, muscle twitches: 0.18 ± 0.06). However, the resulting ROI-based frequency maps differed dramatically between the motion and rest (Fig. 2A). During the motion period, activity was mostly localized to the somatosensory and a lesser extent motor cortex, whereas at rest it was predominantly found in the visual and auditory cortices as well as a rim of lateral cortical areas likely including temporal and parietal cortices.

Noteworthy, the distinct behavioral state-specific activity patterns were highly conserved across experimental animals (Figs. 2-2 and 2B). To quantify this difference, we calculated spatial correlation coefficients (Lewis, 1995) for ROI-based frequency maps recorded during either the same or different behavioral state. First, we compared the maps calculated for three 10-min-long consecutive recordings in the same animal (Fig. 2E) and then compared the median values obtained in 7 mice (Fig. 2F). For all animals tested, spatial correlation coefficients were high for “within the state” comparison and much lower when comparing the maps across the different states. From a statistical point of view (Fig. 2F), the coefficients were highest during the motion state and somewhat lower during the resting state, whereas the comparison between motion and rest produced very small correlation coefficients. The differences between the three groups were statistically significant (One-way Repeated Measure ANOVA F_2, 12_ = 71.1, P < 10^−3^ followed by Holm-Sidak multiple comparisons test; motion vs. rest P = 10^−3^ and P < 10^−3^ for the two remaining comparisons). The almost inverse spatial maps of the local activity during the two behavioral states suggest that cortical networks involved in generation and propagation of this spontaneous activity are tightly controlled in a state-dependent manner.

### Hemispheric asymmetry of the local cortical activity

Next, we used the multi-ROI filters to analyze whether the patterns of coherent spontaneous activity were hemisphere-symmetric (Fig. 3). To do so, for each recording we constructed the total symmetry map (containing only hemisphere-symmetric ROIs) and the semi-symmetry map (containing at least one pair of hemisphere-symmetric ROIs, see Materials and methods) and compared these maps across the animals and the behavioral states. In general, only 6 ± 2.4% of all multi-ROIs showed a stringent hemispheric symmetry (Fig. 3A-C). Another 9 ± 7.3% of multi-ROIs were partially symmetric, whereas 83 ± 6.5 % were asymmetric (the superposition of these ROIs would look similar to Fig. 2A and 2B). The vast majority (73 ± 22.7%) of totally symmetric ROIs was observed during the motion period, and only some 10.3 ± 14% of symmetric activity patterns happened at rest (Fig. 3D-F). Similar results were obtained for semi-symmetry maps, with 75 ± 29% of respective activity patterns observed during the motion and 12 ± 28% observed during the resting period (Fig. 3G-I). Because the cortical activity was more frequent during the motion period (Fig. 2A and 2B), we normalized the number of total/semi-symmetry events during motion/rest to the respective total number of events. The normalized fractions of events remained low (semi-symmetry: 9 ± 2% and 3.4 ± 1% and total symmetry: 6 ± 1 % and 1.5 ± 0.5 % during motion and rest, respectively; mean ± S.E.M). On average, total/semi-symmetry events were 2.3 ± 0.5 times more frequent during motion than during rest. Because at the age studied here the callosal projections, providing interhemispheric communication, are immature (Son et al., 2017; Wang et al., 2007), more frequent occurrence of symmetric events during motion is likely caused by peripheral or subcortical structures.

Subsequently, we asked whether regions contributing to hemisphere-symmetric neuronal activity differ between the two behavioral states. For both total and semi-symmetry maps, the fraction of active pixels amounted to less than 22% of all imaged cortical pixels (Fig. 4A-D).

**Fig. 4.**
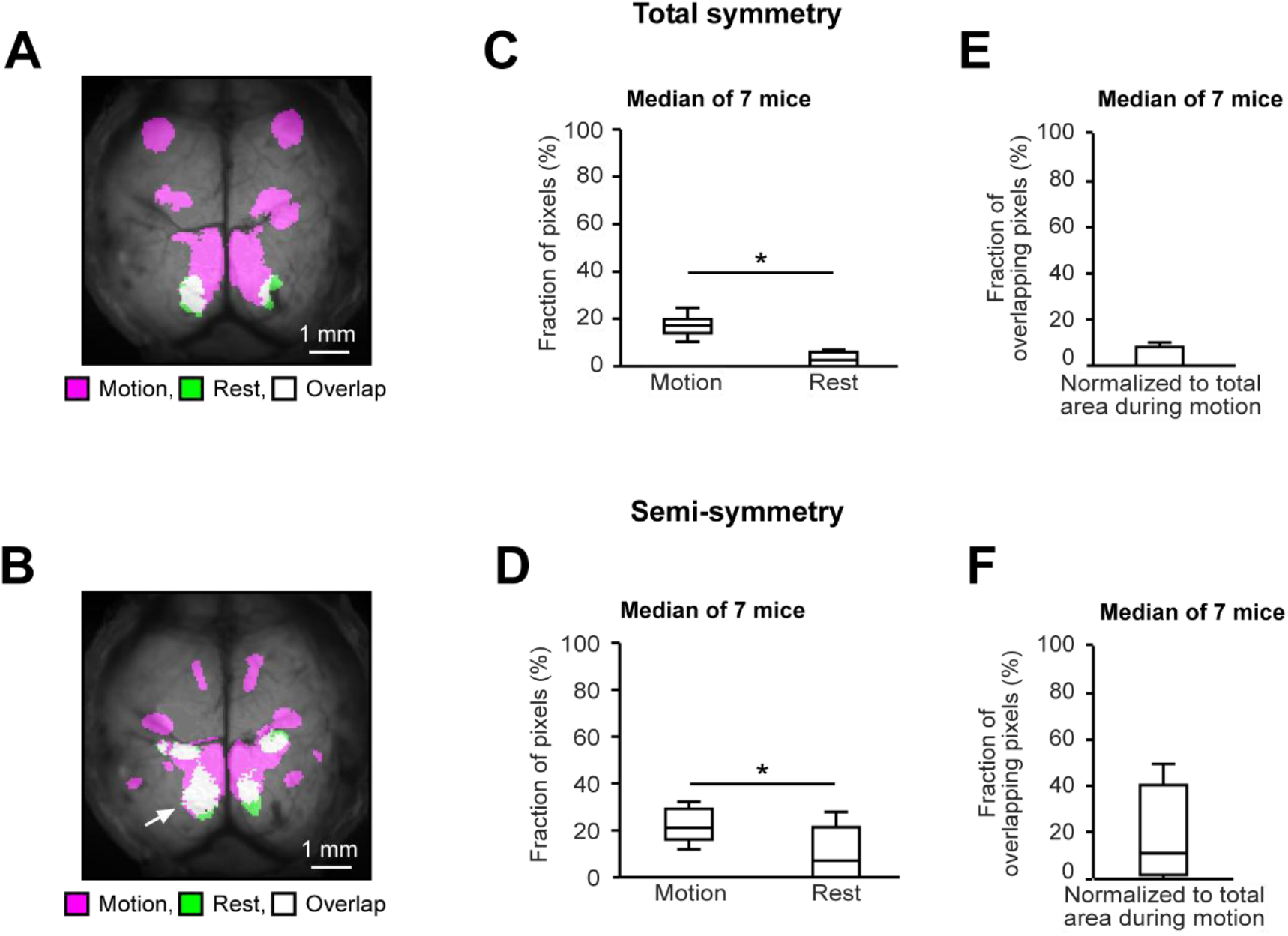
Regions contributing to hemisphere-symmetric cortical activity in the neonatal mouse cortex differ between behavioral states. ***A***, Spatial representation of the overlap between total symmetry maps recorded during motion and resting time periods in a P3 mouse. Purple color indicates the area active during the motion period, green color marks the area active during the resting period and white color shows the overlapping area. ***B***, The same representation as in ***A*** but for the semi-symmetry map obtained in the same experimental animal. The arrow points to the retrosplenial cortex. For display purposes, an experiment with a prominent overlap is shown. ***C***, Box-and-whisker plot showing the number of pixels belonging to the total symmetry map during motion (purple area) and rest (green area), as median (per mouse) fractions of the total number of imaged cortical pixels (n = 7 mice). Obtained values are significantly different (paired Student’s t-test, t_5_ = 7, P < 10^−3^). ***D***, The same analyses as in ***C*** but for the areas contributing to the semi-symmetry map. Obtained values are significantly different (paired Student’s t-test, t_5_ = 5.7, P = 2 × 10^−3^). ***E***, Box-and-whisker plot showing median (per mouse) numbers of pixels belonging to the white area (as in ***A***) normalized to the number of pixels belonging to the purple area (n = 7 mice). ***F***, The same analyses as in ***E*** but for the areas contributing to the semi-symmetry map.

Still, the fraction of pixels active during the motion period (17.2 ± 3.9% and 21 ± 11.1%, for total and semi-symmetry maps, respectively) was significantly higher than the fraction of pixels active at rest (2.5 ± 5.9% and 7 ± 21.3%; Paired student’s t-test; total symmetry: motion vs. rest, t_5_ = 7, P < 10^−3^; semi-symmetry: motion vs. rest, t_5_ = 5.7, P = 2×10^−3^). Although we did identify some anatomical regions involved in totally- and semi-symmetric activities during both behavioral states (the most prominent being the retrosplenial cortex, marked with an arrow in Fig. 4B), in general, the regions involved in these kinds of activity differed between the resting and motion periods. Thus, for the total symmetry maps (Fig. 4E) there were virtually no pixels active both during motion and rest (0 ± 8%). For the semi-symmetry maps, 11 ± 41.8% of pixels active during motion were also active at rest (Fig. 4F).

Taken together, the data clearly documents a profound hemispheric asymmetry of spontaneous neuronal activity at this developmental state. This lack of symmetry is likely due to the immature corpus callosum (Son et al., 2017; Wang et al., 2007), known to coordinate synchronous patterns of activity between hemispheres (McVea et al., 2016). At the same time, we show that in the minority of cases hemisphere-symmetric activity patterns do occur and identify the retrosplenial cortex as a region, frequently involved in such type of activity during both the movement and the resting periods.

### Behavioral state-specific maps of simultaneously active cortical subregions

Multi-ROI filters, automatically provided by a nonnegative matrix factorization algorithm, contain synchronously active areas. We used this property to visualize brain regions active simultaneously during either the movement or the resting states. First, we determined the anatomical location of all multi-ROI subregions using the map similar to the one shown in Fig. 1A but including also the retrosplenial cortex (Fig. 5-1A). Subsequently, we constructed the state-dependent maps (see Materials and methods for details), in which each edge connected brain regions, which were consistently simultaneously active in a behavioral state-specific manner within and across experimental animals (Fig. 5-1B-D). The circles indicate instances in which pairs of active subregions of a multi-ROI were located within the same anatomical region.

In both behavioral states, the simultaneously active subregions were found either close to each other (e.g., within the same anatomical region) or up to several mm apart, as exemplified in Fig. 5-1B-D. In general, the center-to-center distances between simultaneously active subregions ranged from 0.3 to 5.8 mm (1^st^ - 99^th^ percentile), with a median of 2.1 ± 0.4 mm. When comparing median (per mouse) distances between the subregions, active during the motion and the resting states, no significant difference was found (motion: 2.1 ± 0.4 mm, rest: 2.2 ± 0.2 mm; Wilcoxon signed-rank test z = 0.52, P = 0.69, n=6 mice; Fig. 5-2B). However, consistent with the general activity pattern described in Fig. 2, the regions contributing to the simultaneous activity map differed dramatically between the motion and the resting state. During the motion state, the synchronously active subregions were mostly located in the anterior (motor, somatosensory) cortical regions, shifting to the posterior (e.g., visual) cortical regions during the resting state (Fig. 5-1C and 5-1D). Interestingly, auditory and retrosplenial cortices contributed to the map of simultaneous activity during both behavioral states. During the motion period, however, the retrosplenial cortex was predominantly active together with the motor cortex, while during the resting state it was predominantly active together with the visual cortex. The median simultaneous activity map obtained for all recorded mice (n = 6; Fig. 5-1D) assured the solidity of the described above findings.

### Waves of activity propagating through the neonatal cortex

So far when describing the observed activity patterns we have mostly considered their spatial dimension. With the temporal dimension added (see Materials and methods for details), the observed neonatal activity patterns turned into waves of activity either propagating in a given direction (e.g. Fig. 5A, left panel) or waxing and waning in magnitude and spatial area involved without any clear movement of the wave’s center of mass (so-called stationary waves). Of all propagating waves observed, the majority (81 ± 25%) were recorded during the motion period with a significantly smaller fraction of waves (19 ± 25%; Paired student’s t-test, t_5_ = 3.2, P = 0.02) being recorded at rest (Fig. 5B, upper panel). A similar trend was also observed for stationary waves (motion: 81 ± 36%, rest: 19 ± 34%) but the difference did not reach the level of statistical significance (Fig. 5B, lower panel).

Next, we characterized the fractions of waves invading different anatomical regions (i.e. motor, somatosensory, auditory, visual, and retrosplenial cortices) of the right and the left hemispheres during the two behavioral states. For both propagating and stationary waves we did not observe any differences between the left and the right hemisphere for any condition tested (Repeated Measures ANOVA; Fig. 5-3A: F_1, 5_= 3.5, p=0.12; Fig. 5-3B: F_1, 5_= 2.8×10^−4^, p=0.98; Fig. 5-3C: F_1, 5_= 4.1, p=0.1; Fig. 5-3D: F_1, 5_= 10^−3^, p=0.98). Therefore, for further analyses, we averaged the data from both hemispheres (Fig. 5C-F). During the motion state the highest fraction of both propagating (23 ± 4.5%, Fig. 5C) and stationary (27 ± 8.5%, Fig. 5D) waves invaded the somatosensory cortex, whereas at rest the waves mostly invaded somatosensory, auditory, and visual cortices (9.8 ± 4%, 15.8 ± 4.5%, 11.8 ± 6% for propagating and 13 ± 9.5%, 18.8 ± 7.5%, 9.8 ± 4.5% for stationary waves, respectively).

As to the wave frequency (Fig. 5-4A and C), it was highest in the somatosensory cortex (2.3 ± 0.45 waves/min for propagating and 2.7 ± 0.85 waves/min for stationary waves) during motion and in the auditory, visual and somatosensory cortices (propagating: 1.58 ± 0.45, 1.18 ± 0.6, 0.98 ± 0.4 waves/min; stationary: 1.88 ± 0.75, 0.98 ± 0.45, 1.3 ± 0.95, waves/min, respectively) during rest. A similar frequency range was reported previously for different types of early intrinsic cortical activity in rats and mice (Adelsberger et al., 2005; An et al., 2014; Brockmann et al., 2011; Hanganu et al., 2006; Kirmse et al., 2015). To account for different sizes of the cortical areas under study, in the next step we normalized fractions of waves, invading different cortical regions, to the area of that particular region (Fig 5-5A and C). This analysis stressed the contributions of the somatosensory cortex to the activity during the motion and that of the auditory and visual cortices to the activity during the resting state.

Next, we analyzed locations of pacemakers (defined as described in Materials and methods), initiating a given type of activity. During the motion state, the pacemakers were mostly located in the somatosensory cortex (25.5 ± 6.5% of pacemakers for propagating (Fig. 5E) and 27.3 ± 8.5% for stationary (Fig. 5F) waves, respectively). During the resting state the pacemakers were mostly distributed between the somatosensory, auditory, and visual cortices (11.8 ± 5.5%, 18 ± 5%, 13.5 ± 7.5% of pacemakers for propagating and 13 ± 10%, 19 ± 7.5%, 9.8 ± 5% for stationary waves, respectively). The frequency, with which pacemakers appeared in different cortical regions (Fig. 5-4B and D), was highest (2.55 ± 0.65 per min for propagating and 2.73 ± 0.85 per min for stationary waves) in the somatosensory cortex during motion and in the auditory, visual and somatosensory cortices (propagating: 1.8 ± 0.5, 1.35 ± 0.75, 1.18 ± 0.55 per min; stationary: 1.9 ± 0.75, 0.98 ± 0.5, 1.3 ± 1, per min, respectively) during rest.

Normalization to the relative size of the cortical region showed that both for propagating and stationary waves the pacemaker density was highest in the somatosensory cortex during motion and in the auditory, visual and retrosplenial cortices during rest (Fig. 5-5B and D). The overall similarity in the distributions of waves, invading a given cortical area, and their pacemakers suggests that the majority of waves spread over rather short distances.

To test this assumption, we analyzed the distances traveled by the centers of mass of all propagating waves in our dataset. In general, the propagating waves spread over distances of 0.22 - 4.8 mm (1^st^ – 99^th^ percentile), with the median of 0.47 ± 0.02 mm (n = 6 mice, not differentiating between motion and rest). The apparent speed of the center of mass ranged from 0.06 to 3.5 mm/s (1^st^ – 99^th^ percentile), with the median of 0.38 ± 0.05 mm/s (n = 6 mice). When comparing the properties of waves propagating during the motion and the resting states (Fig. 5-6), there was no significant difference between the distributions of median (per mouse) propagation distances (motion: 0.47 ± 0.02 mm, rest: 0.47 ± 0.06 mm; Paired student’s t-test t_5_ = 0.34, P = 0.75, n = 6 mice; Fig. 5-6B) but the respective propagation speed (Fig. 5-6D) was significantly different (motion: 0.42 ± 0.05 mm/s, rest: 0.28 ± 0.13 mm/s; Paired student’s t-test t_5_ = 4.3, P = 0.01, n = 6 mice).

Thus, the spontaneous neuronal activity in the dorsal cortex of neonates contains a rich and spatially complex pattern of both stationary and propagating waves. The majority of the propagating waves are associated with the animal’s movement, whereas stationary waves seem to occur with roughly similar incidence during the movement and the resting periods. Interestingly, the waves of both types are triggered predominantly in the somatosensory or auditory cortices.

### Functional connectivity map of early cortical activity

Spontaneous correlated neuronal activity is believed to represent a functional template for activity-dependent maturation of intracortical connections and the refinement of functional units underlying information processing at adulthood (Hanganu-Opatz, 2010; Luhmann et al., 2016). To test for functional connectivity between the different cortical regions (Fig. 6), we used a sparse inverse covariance matrix estimation algorithm (see Materials and methods for details).

**Fig. 6.**
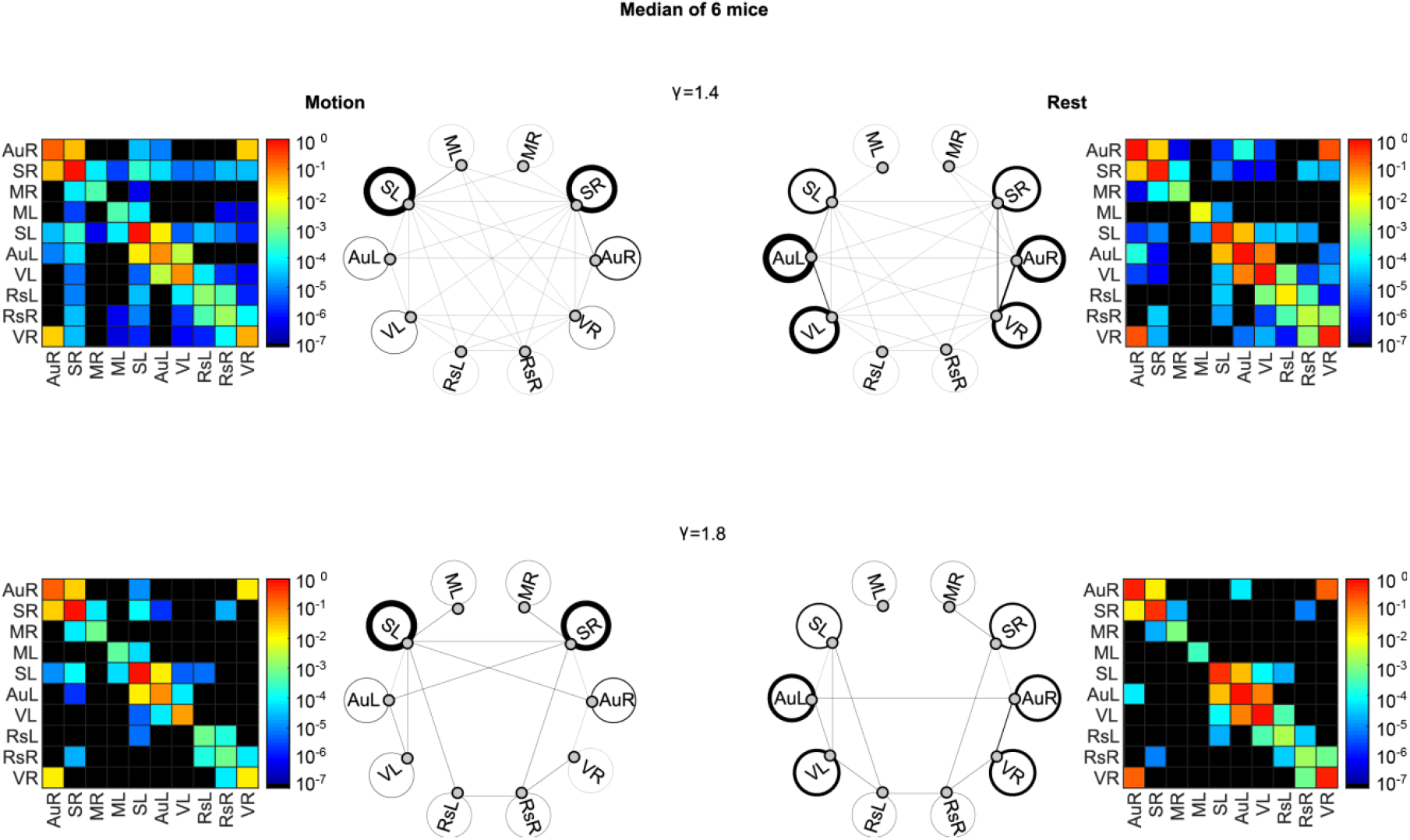
Functional connectivity map. Direct connectivity maps calculated for motion (left panels) and resting (right panels) periods with two different values of γ (see Materials and methods). Each connectivity map is shown in the matrix as well as trivial graph format. In the latter format, nodes represent cortical regions of interest predefined in Fig. 5-1A, and edges between the two nodes show the direct connectivity between the two cortical regions. Edge thickness represents the median strength of connections between the two cortical regions. Similarly, circles depict connectivity within the given cortical region, and the thickness of each circled line shows the median strength of connections in this cortical region. Note that in the matrix format the strength of connectivity is shown on a logarythmic scale, providing a higher dynamic range. Median of data obtained from 6 different animals. Color-coded scale bars show median connectivity strength within and among cortical regions (see Materials and methods for details).

This analysis was shown to measure the direct association between the two brain regions removing the contributions caused by global or third-party effects (for more details on the method we refer the reader to ref. (Huang et al., 2010)).

The level of sparseness of the algorithm’s outcome was controlled by the tuning parameter γ. Thanks to the monotone property of the algorithm used (Huang et al., 2010), increasing γ monotonically increases the level of sparseness of cortical connectivity at the expense of weaker connections. As, shown in Fig. 6-1, at γ=1.4 the sparsity of the connectivity map was reached for the first time for both the motion and resting states. Beyond γ=1.8 the number of connections was strongly diminished. At γ=1.4 we have observed direct functional connections within and in-between of many anatomical regions (Fig. 6, upper panel), defined as illustrated in Fig. 5-1A. However, the majority of these connections were weak (Fig. 6, upper panel) and disappeared with an increasing γ (Fig. 6, lower panel). The remaining strong connections emphasized the short-range intraregional connectivity as well as behavioral state-specific long-range connections between the more anterior regions during motion. Interestingly, the method identified a strong and direct functional link between the ipsilateral retrosplenial and somatosensory cortices, present during both behavioral states, as well as some prominent behavioral state-specific connections (Fig. 6).

Together, these data reveal that in neonatal mice functionally connected cortical subregions are distributed through the entire dorsal cortical surface. The connectivity pattern is characterized by dense short-range connections, linking together different subregions of the given anatomical area and immediately adjacent cortical areas, as well as sparse long-range connections. The latter differ substantially between the resting and the motion state. These differences are highly conserved across experimental animals thus reflecting typical activity patterns of the neonatal brain.

## Discussion

The current study reveals behavioral state-specific large-scale cortical maps in the dorsal cortex of neonates (Fig. 2). The motion- and rest-associated maps are almost orthogonal in nature and are built of patches of recurrent endogenous activity. The existence of these reciprocal maps could not have been anticipated based on the previously available data (Adelsberger et al., 2005; An et al., 2014; Anton-Bolanos et al., 2019; Golshani et al., 2009; Hanganu et al., 2006; Khazipov et al., 2004; McVea et al., 2012; Smith et al., 2018; Valeeva et al., 2019). For both behavioral states studied, the recurrent endogenous activity showed little hemispheric symmetry and consisted of a complex mixture of (i) small local events, often limited to a given anatomical region, (ii) larger events, connecting the neighboring cortical regions, as well as (iii) simultaneous or correlated activity patterns, binding together activities of different cortical regions located up to 6 mm apart. Moreover, the patterns of the binding activity were different between the different behavioral states with somatosensory-motor areas functionally connecting during the motion and the auditory-visual areas functionally connecting during the rest.

Interestingly, frontal and parieto-occipital activity patterns were revealed (by EEG recordings) also in the perinatal human cortex (Omidvarnia et al., 2014). The latter turned out to be vigilant state-specific with parieto-occipital activity patterns observed during the active and a more widespread increase in connectivity throughout the cortex with notable involvement of the frontal areas observed during the quiet sleep (Tokariev et al., 2019). Despite the robust neuronal activity, the accompanying local hyperemia is apparently absent in both perinatal humans (Omidvarnia et al., 2014) and mice (Movie 2-1, (Kozberg et al., 2016)), disabling the analyses of this activity by fMRI. The “small-worldness” represents another feature, similar for both species (Figs. 6 and (Omidvarnia et al., 2014)). Compared to adults and full-term newborns, activity patterns in the perinatal, especially preterm, human cortex are also known for the reduced interhemispheric synchrony and symmetry (see (Koolen et al., 2014; Kwon et al., 2015; Meijer et al., 2016; O’Toole et al., 2019) and references therein). In both mice and humans, this is likely due to the immature corpus callosum, developing in humans between 35 and 37 weeks of gestation (Meijer et al., 2016; Son et al., 2017; Wang et al., 2007).

Our work has also identified the unique binding role of the retrosplenial cortex, an area (i) involved in both motion- and rest-related ongoing activities, especially in their hemisphere-symmetric subtypes, and (ii) maintaining strong long-range functional connections with many other studied cortical regions. Although the consensus on the precise function of this cortical area is still missing, evidence from both human and animal studies points to its role in spatial navigation, visuospatial integration, and hippocampus-related learning and memory (Czajkowski et al., 2014; Vann et al., 2009). Moreover, in adult humans and mice retrosplenial cortex was shown to be active during the resting brain state, thus belonging to the so-called “default mode network” (Chan et al., 2015; Vann et al., 2009). Whereas many areas of the default mode network are known to decrease their activity upon engagement into a cognitive task (Anticevic et al., 2012), this is not the case for the retrosplenial cortex, which increases its activity, for example, during spatial navigation or autobiographical memory retrieval (Vann et al., 2009). Of special interest is also the fact that although areas involved in the default mode network vary across different age groups, this is not the case for the retrosplenial cortex, which represents its inherent part from early infancy through adolescence into adulthood (Vann et al., 2009). We hypothesize that both in humans and now also in mice from early on the retrosplenial cortex represents a functional bridge between the default mode network active at rest and the task- or movement-specific neural networks coordinating ongoing activity in the sensorimotor system. To do so, the retrosplenial cortex engages in a behavioral state-specific coherent activity with the respective cortical areas: visual cortex at rest and somatomotor cortices during motion.

The long-range nature of the latter functional connections is in marked contrast with the developmental state of the immature P3 mouse brain, in which long-range anatomical connections are immature (Hartung et al., 2016; Tagawa and Hirano, 2012). Interestingly, a recent study of the ferret visual cortex also discovered long-range (albeit intra-regional) functional connections at the developmental stage (P21 - 22 in ferret) similar to the one studied here (Smith et al., 2018). To explain their data the authors presented a dynamical rate network model, which in a regime of strongly heterogeneous local connectivity and moderate input modulation produces pronounced long-range correlations, similar to the ones observed in an experiment. Noteworthy, the spatial structure of the correlated activity produced by the model was fairly robust against changes in the input drive strength, in agreement with the author’s data that long-range correlations persisted in the immature cortex even after silencing the main driving input (i.e. spontaneous retinal waves reaching the visual cortex via the lateral geniculate nucleus (Smith et al., 2018)). The described above findings suggest that the neonatal brain utilizes local spontaneously active inputs to drive the correlated activity of distant cortical regions, thus providing the template for activity- and Ca^2+^-dependent growth and branching of long-range axonal projections (Tagawa and Hirano, 2012). Indeed, anatomical studies in adult rodents suggest that the retrosplenial cortex, to stay with this example, is directly linked with anterior cingulate, motor, and visual cortices (Oh et al., 2014; Vann et al., 2009; Yamawaki et al., 2016).

Local self-initiated activity and associated spatially-restricted Ca^2+^ signals represent the predominant type of activity also in our study. In-depth characterization of this type of activity revealed its rich spatiotemporal structure comprising distributed patches of coherent activity in spatial as well as propagating and stationary waves in spatiotemporal domains. Strikingly, local cortical activity patterns recorded during the motion and the resting states populated different brain areas. During the motion, activity was restricted to the somatosensory-motor area (compare active area in Fig. 2B to the map of the adult mouse brain in ref. (Vanni et al., 2017)), consistent with the cumulative knowledge derived from earlier reports (An et al., 2014; Khazipov et al., 2004; McVea et al., 2012; Tiriac et al., 2012) as well as with recent data showing that active wake movements suppress spontaneous neural activity in the visual cortex of neonatal rats (Mukherjee et al., 2017). In contrast, during the rest local activity spared the above areas being mostly restricted to the visual, auditory, and retrosplenial cortex as well as lateral cortical areas, such as, for example, the temporal cortex, known for its movement-independent spontaneous network activity (Adelsberger et al., 2005). In-depth analyses of the propagating and stationary waves showed that the local activity has the highest frequency in the somatosensory cortex during motion and the highest density in the auditory cortex during rest (Figs. 5-4 and 5-5).

The described above segregation of the activity patterns is surprising also in view of the fact that during the developmental stage studied here both motion and rest are behaviorally heterogeneous. Indeed, motion comprises both generalized movements happening during wakefulness and muscle twitches, mainly observed during active sleep (Seelke et al., 2005; Tiriac et al., 2014). Similarly, the resting period might also include both quiet wakefulness and quiet sleep. Because under our experimental conditions the overall duration of muscle twitches comprised only 5 - 16% of the total movement time and the amount of data variance related to twitches was only 0.2 ± 0.6%, the described here motion-related activity pattern is dominated by the one caused by wake-related generalized movements. Still, it is consistent with the activity pattern, seen by others during spontaneous muscle twitches (McVea et al., 2012; Tiriac et al., 2012). Thus, under conditions when animal’s limbs are free to interact with each other and the supporting surface, both kinds of self-generated movements discussed above (predominantly wake-related generalized movements: current study; muscle twitches refs. (Khazipov et al., 2004; McVea et al., 2012; Tiriac et al., 2014; Tiriac et al., 2012)) cause a robust and predominantly reafferent neuronal activity in the somatosensory-motor area of the cortex. However, the measured here high amount of data variance associated with self-initiated generalized movements is in contrast to the belief that “reafference from twitches triggers cascades of neural activity throughout the sensorimotor system at ages when wake-related movements largely fail to do so” (Del Rio-Bermudez et al., 2020).

## Acknowledgments

We thank E. Zirdum, A. Weible, and K. Schoentag for technical assistance and Matthias Bethge for input concerning procedures for data analyses. This work was supported by DFG Grant GA 654/13-1(belonging to the Research Unit FOR 2715), to O.G.

## Author contributions

O.G. conceived the study. Y.K. and O.G. designed and performed the experiments. N.M. and A.B. developed analysis procedures and N.M. analyzed the data. O.G., N.M., and Y.K. wrote the manuscript. All authors commented on the manuscript.

## Data and software availability

The data and the Matlab code used are available from the corresponding author upon reasonable request.

