## Extended Data for "Stable behavioral state-specific large scale activity patterns in the developing cortex of neonates"

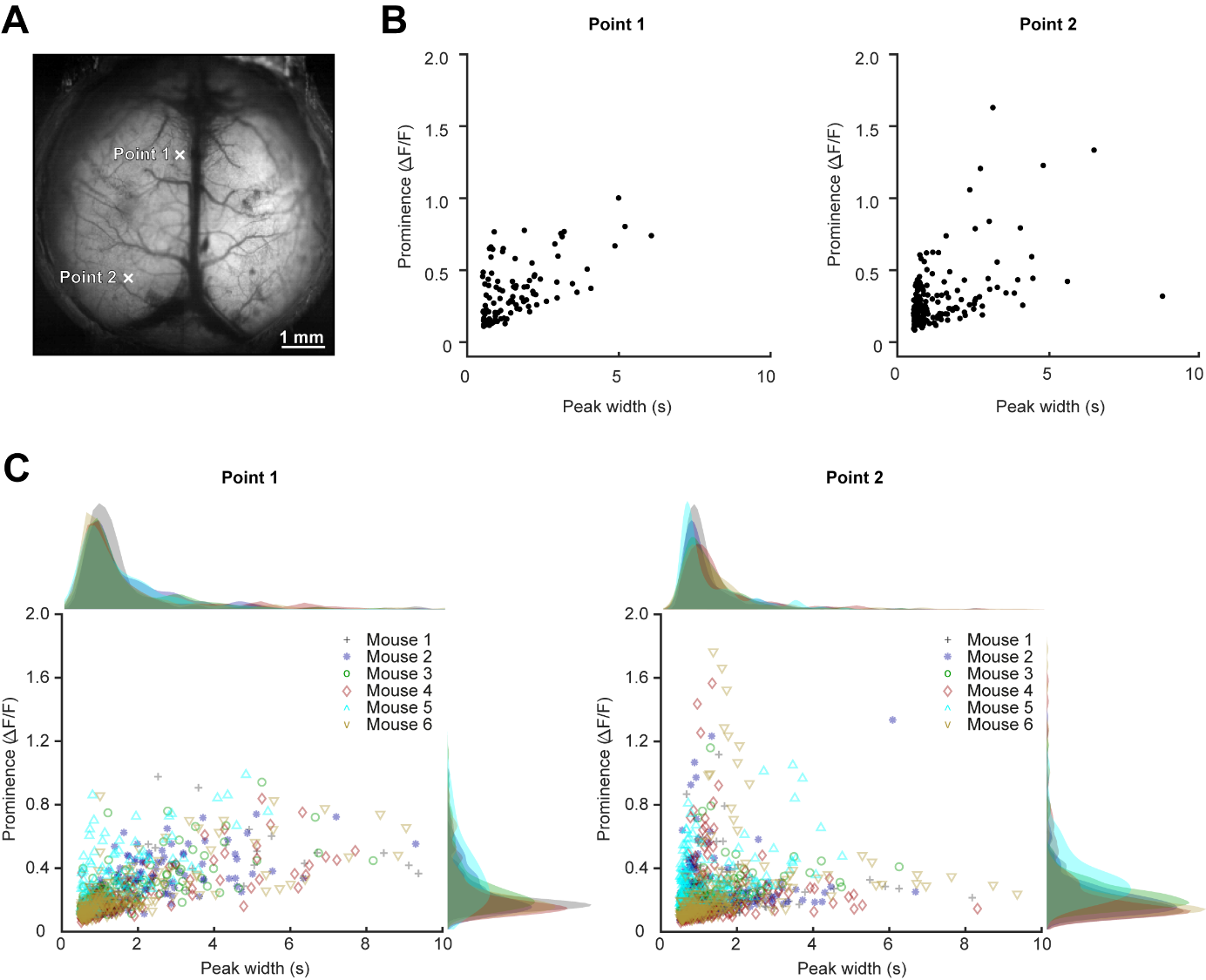

**Figure 1-1. Activity patterns observed in nestin-Cre x Ai95 (RCL-GCaMP6f)-D mice.** ***A***, Top view on a P3 mouse cortex taken through an intact skull. Here and in ***B*** the data are from mouse 5 (Fig. 2-2). White crosses indicate centers of squares (5 x 5 pixels) used to extract the mean fluorescence traces for frontal (Point 1) and visual (Point 2) cortex recordings analyzed in ***B***, ***C***. ***B***, Scatter plot of prominence versus full width in half magnitude (FWHM) values, calculated using Matlab's peak detection function, for peaks detected in fluorescence traces recorded in the frontal (Point 1) and visual (Point 2) cortices. Scatter data shown in ***B*** come from three 10-min-long recordings (same mouse as shown in ***A***). ***C***, Scatter-histograms of prominence versus FWHM values for traces recorded in the frontal (left) and the visual (right) cortices of 6 mice. Data from each mouse is shown with a different symbol. Kernel densities of peak's width values are plotted above x-axis and those of peak's prominence values are plotted on the right side of y-axis (same color coding as in the scatter plot).

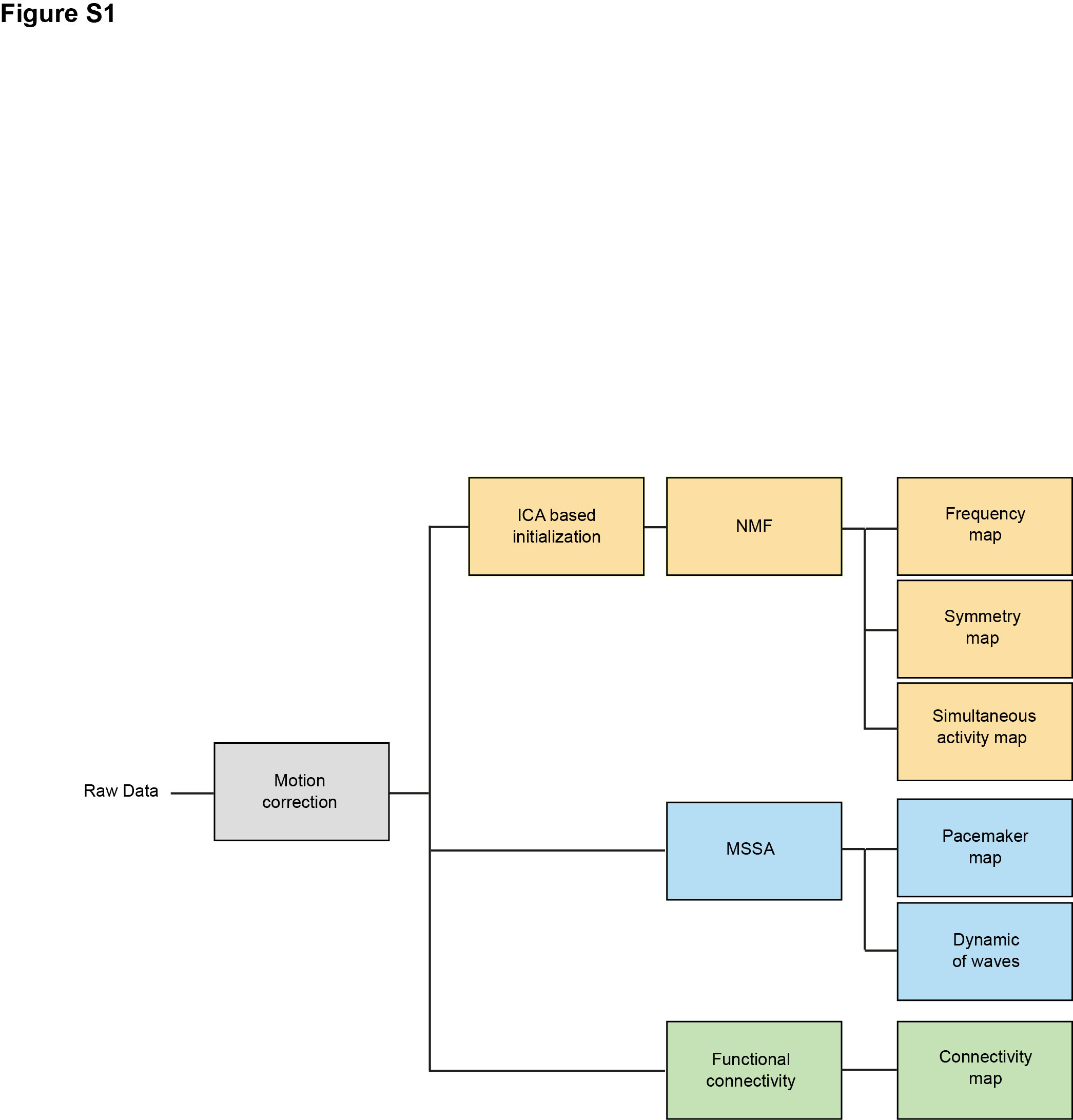

**Figure 2-1. Analyses pipeline.** Schematic representation of the analyses pipeline, explained in the method section. The preprocessing step (motion correction) and 3 branches of analyses are shown in different colors. Matlab (R2016b) image registration toolbox was used for motion correction. Independent Component Analysis (ICA) served as an initialization method for Non-negative Matrix Factorization (NMF). Multi-channel Singular Spectrum Analysis (MSSA) was applied to study dynamical behavior of waves and locations of their pacemakers.

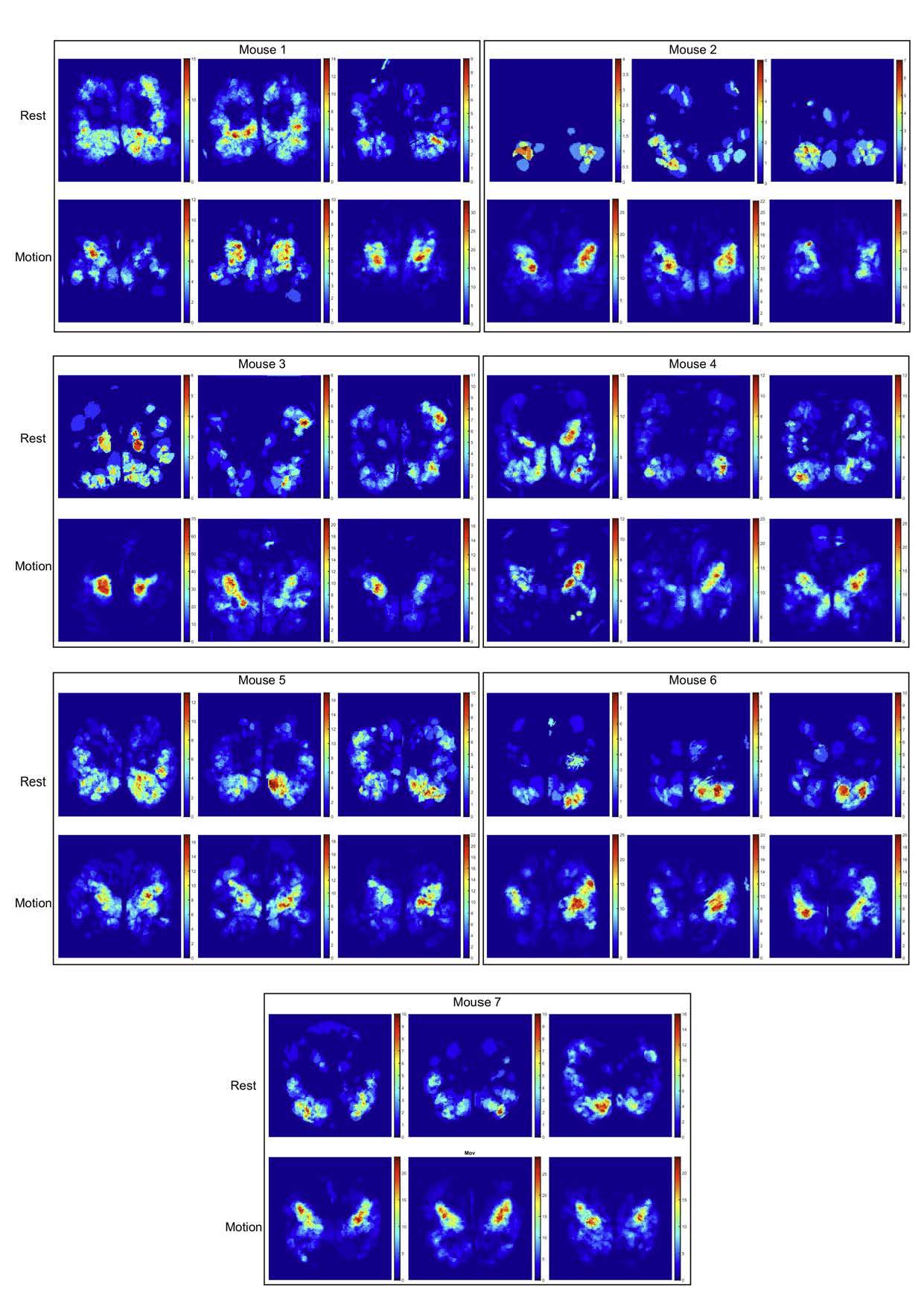

**Figure 2-2. ROI-based frequency maps of local activity**. Frequency maps obtained during motion and rest for all mice used in our experiments. Each box contains two rows to separate maps during rest and motion and 3 columns showing the data from 3 consecutive 10 min recordings. Scale bars show number of events in t=10 min.

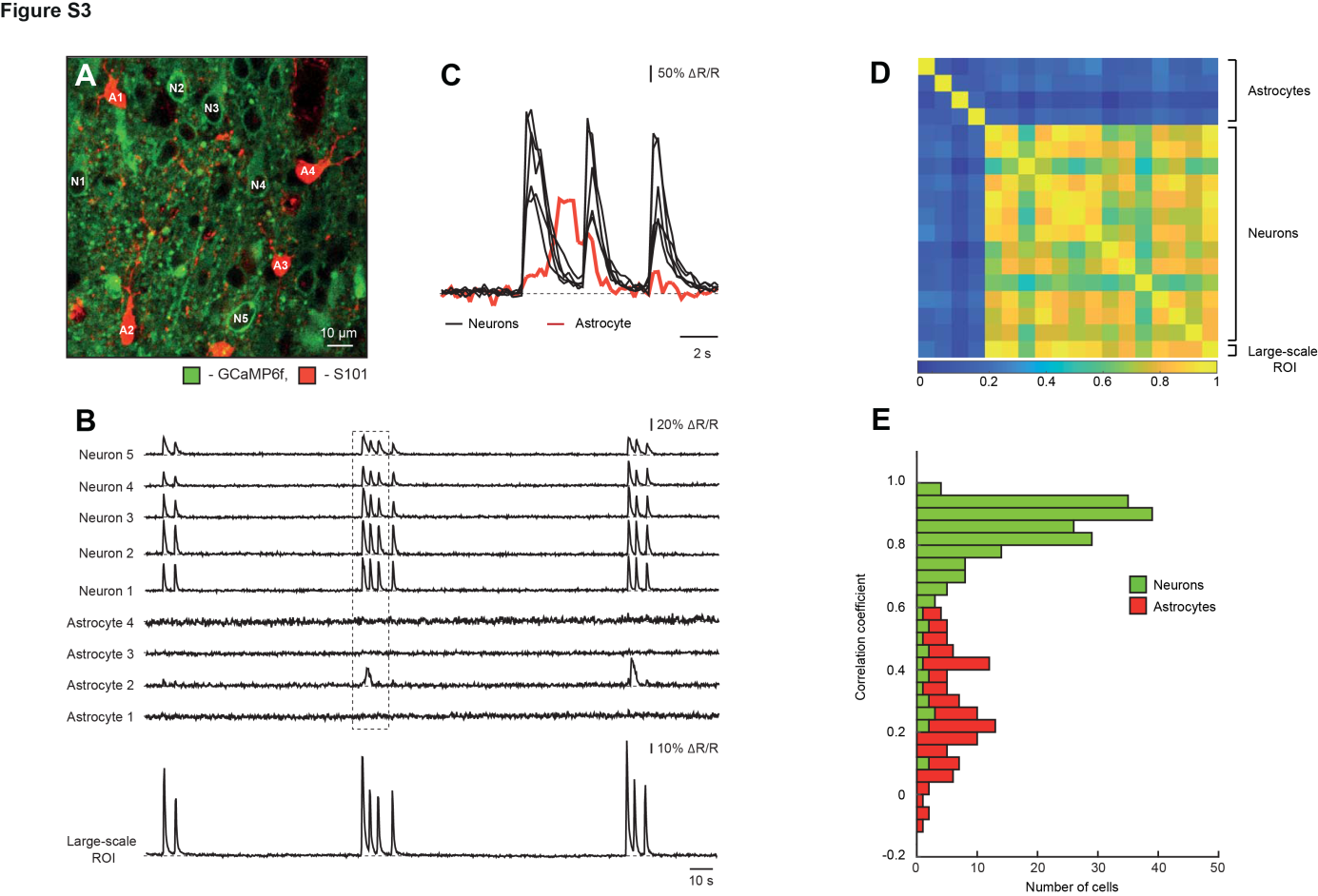

**Figure 2-3. The nature of large-scale cortical activity.** ***A***, An image of neurons labeled with GCaMP6f (green) and astrocytes labeled with both GCaMP6f and S101 (red). Note that for analyses we chose the fields of view containing relatively high density of astrocytes. ***B***, Spontaneous Ca^2+^ signals recorded from individual neurons and astrocytes, marked with respective symbols (i.e. N1-N5 and A1-A4) in ***A***. The lower trace represents the averaged Ca^2+^ signal recorded from the entire field of view shown in ***A***. ***C***, Expended view on the data shown within the boxed region in ***B***. Overlapping neuronal signals (Neurons 1-5) are shown in black. The red trace belongs to Astrocyte 2. ***D***, A correlation matrix plot illustrating correlation coefficients between the traces obtained from 4 astrocytes shown in ***A***, all neurons that can be unequivocally identified in *A* and the respective large-scale ROI. ***E***, Histogram illustrating correlation coefficients between all neuron/respective large scale ROI (green, n=190) and astrocyte/respective large scale ROI (red, n=119) pairs analyzed in this study.

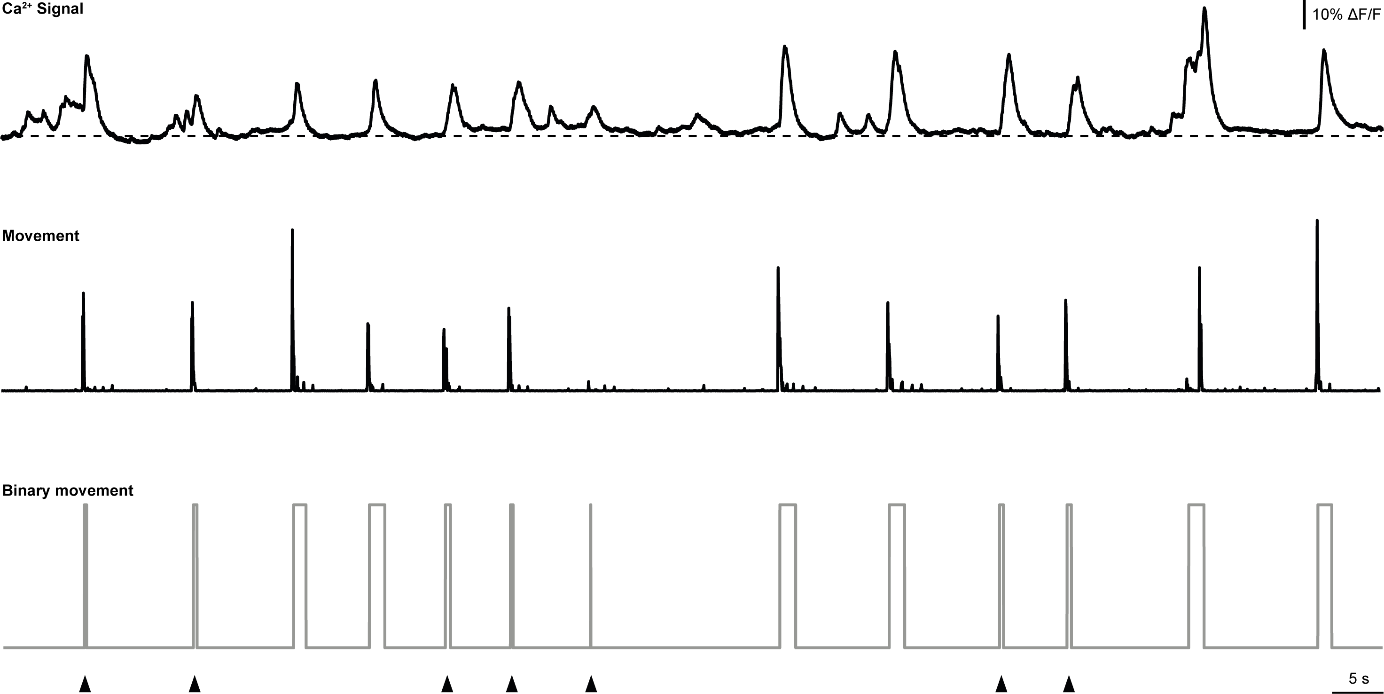

**Figure 2-4. Example traces illustrating Ca^2+^ signals recorded in the somatosensory cortex and corresponding animal movements.** A representative ΔF/F trace recorded from the SR area delineated in Fig. 1A (upper) and the respective raw (middle) as well as binary (lower) animal movement signals, generated as described in Materials and methods. Arrowheads mark muscle twitches. Same experiment as the one shown in Fig. 1*A*-*C*.

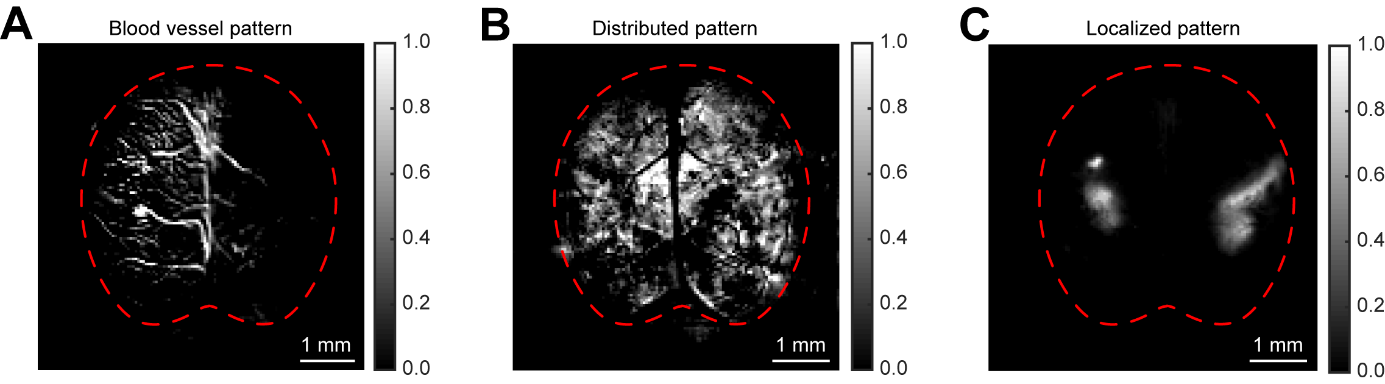

**Figure 2-5. Patterns of activity extracted by the NMF algorithm.** Representative examples of spatial filters containing ***A***, blood vessels pattern, ***B***, distributed changes in fluorescence contaminated by the blood vessel pattern and ***C***, groups of active neighboring pixels, clearly discernible from the background. For display purpose, in all three panels the image intensity is scaled to fit the range from 0 to 1. The red dashed line delineates the borders of the recorded cortical area. Data shown are from mouse 1 (Fig. 2-2).

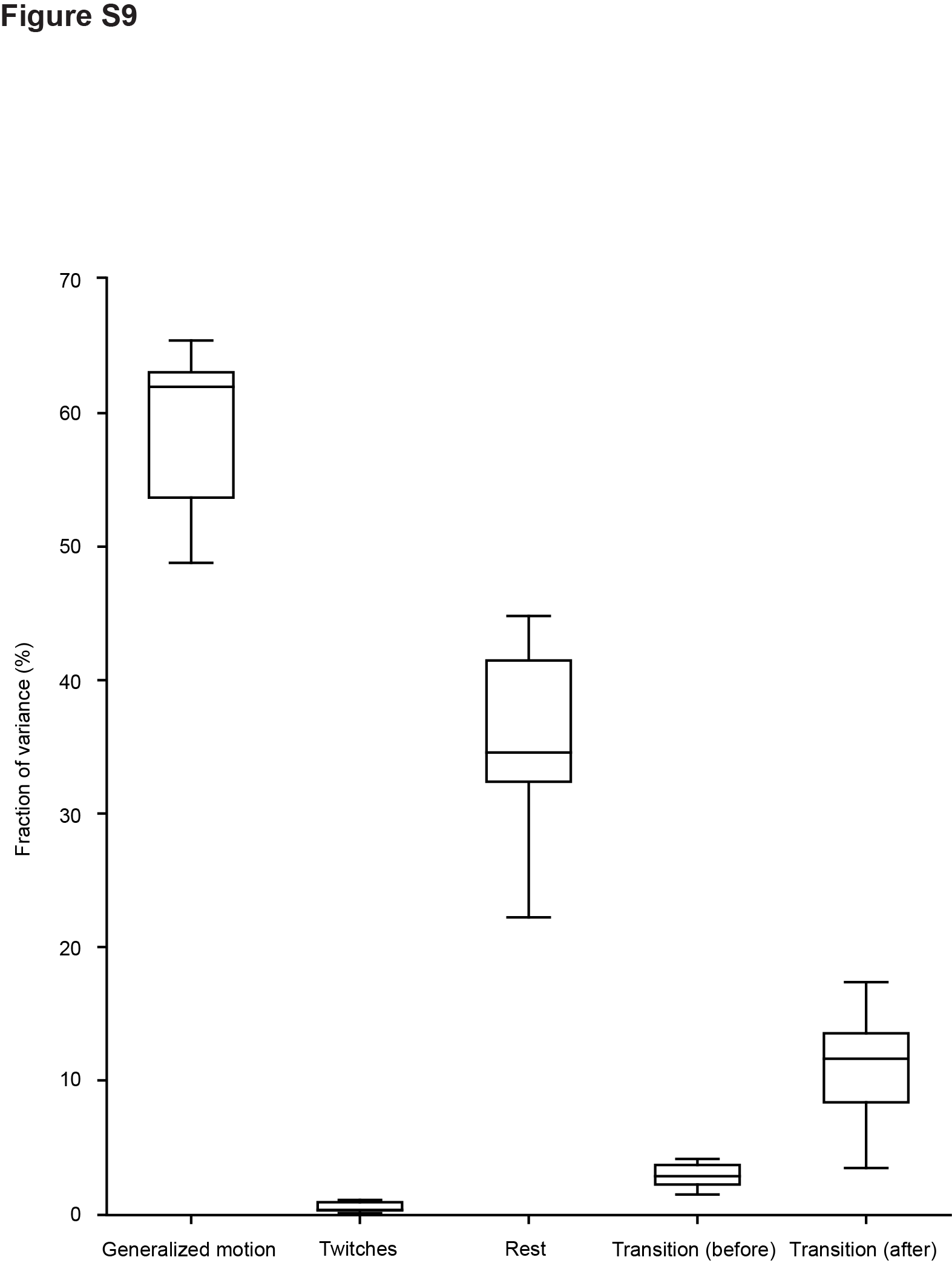

**Figure 2-6. Explained variance in different animal states.** Box-and-whisker plots showing fractions of variance in imaging data related to each behavioral state (n=7 mice).

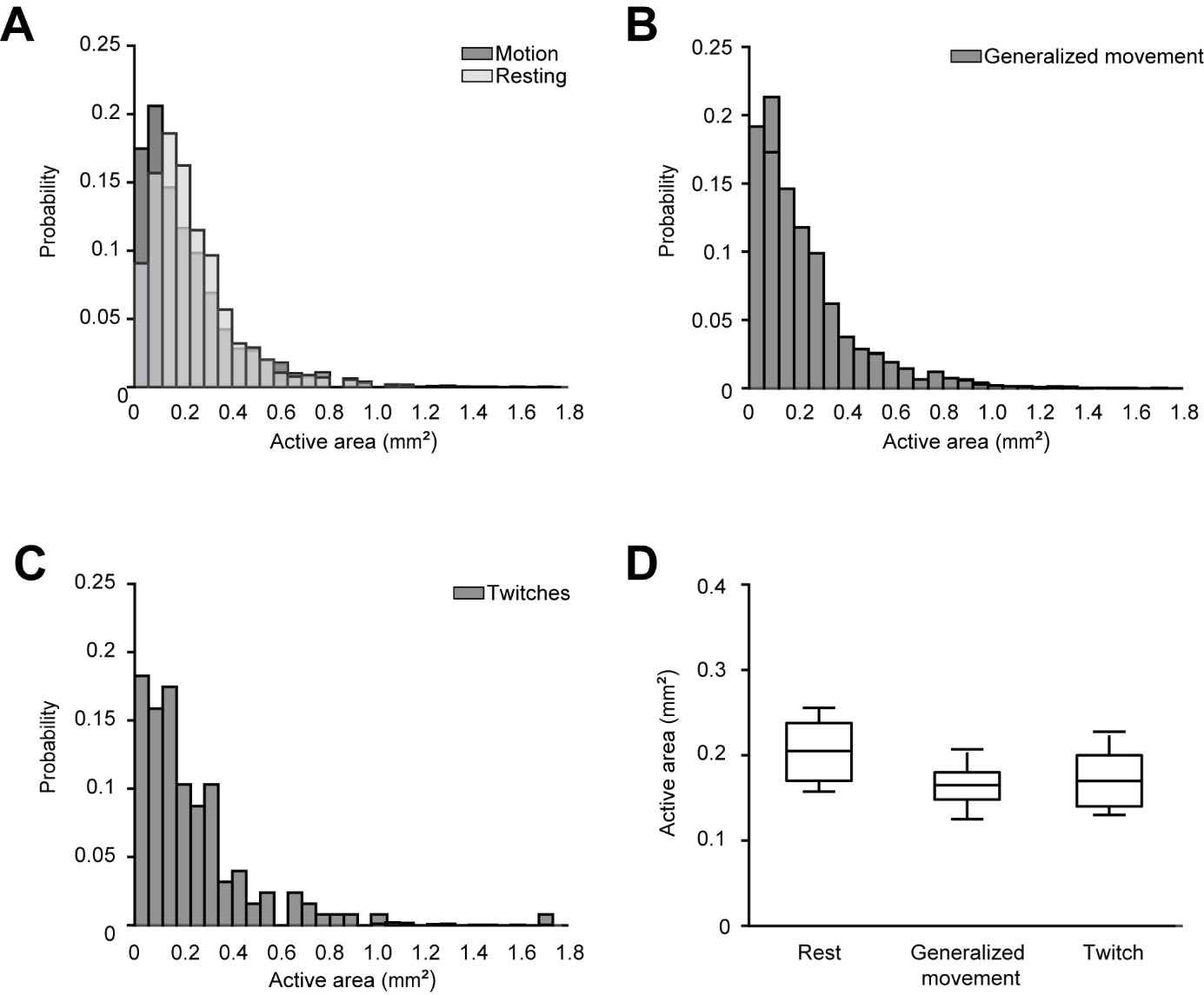

**Figure 2-7. Sizes of coherently active areas contributing to local cortical activity.** ***A***, Normalized histograms of the size of areas active during motion and resting time periods, respectively. ***B***, Normalized histogram of the size of areas active during generalized movements. ***C***, Normalized histogram of the size of areas active during muscle twitches. ***D***, Box-and-whisker plot showing median (per mouse) sizes of areas active during the three above mentioned states (in total n = 8621 areas from 6 mice). As shown by the One-way Repeated Measure ANOVA (F_1.13*, 5.65*_ = 3.8, P = 0.1, *denotes Greenhaus-Geisser sphericity correction), the sizes are not significantly different.

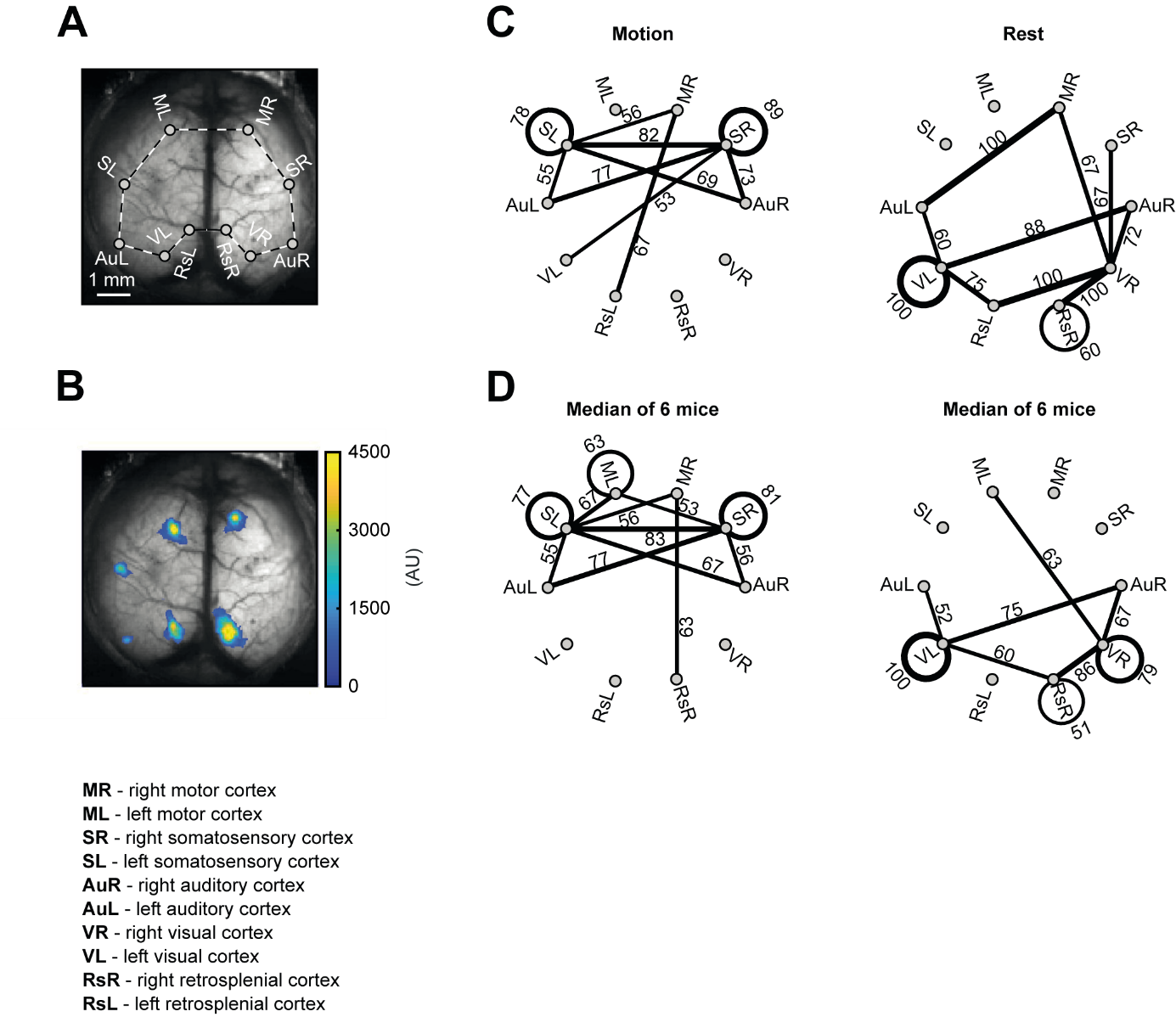

**Figure 5-1. Map of simultaneously active cortical regions.** ***A***, Top view on a P3 mouse cortex. Dots are positioned within the respective cortical areas of interest, listed under the image. ***B***, Representative active subregions belonging to one multi-ROI filter, projected on a gray scale image of a P3 mouse cortex (mouse 5, Fig. 2-2). Fluorescence signals are color-coded with warmer colors indicating higher signal intensity. ***C***, Representative multi-ROI-based maps of simultaneously active regions recorded during the two different behavioral states: motion and rest (3 consecutive 10-min-long image series). Nodes represent cortical areas pre-defined in *A* and edges between two nodes depict cases when two simultaneously active subregions were located in the two cortical areas. Similarly, circles depict cases when two simultaneously active subregions were located in the same cortical area. Numbers along the edges (or circles) represent the normalized number of transients, i.e. the fractions of cases occurring either during the motion (left) or the rest (right), respectively. Here and in ***D*** only the edges and circles with values of ≥50% are shown. ***D***, Same analyses as in ***C*** but illustrating median data obtained in 6 different mice.

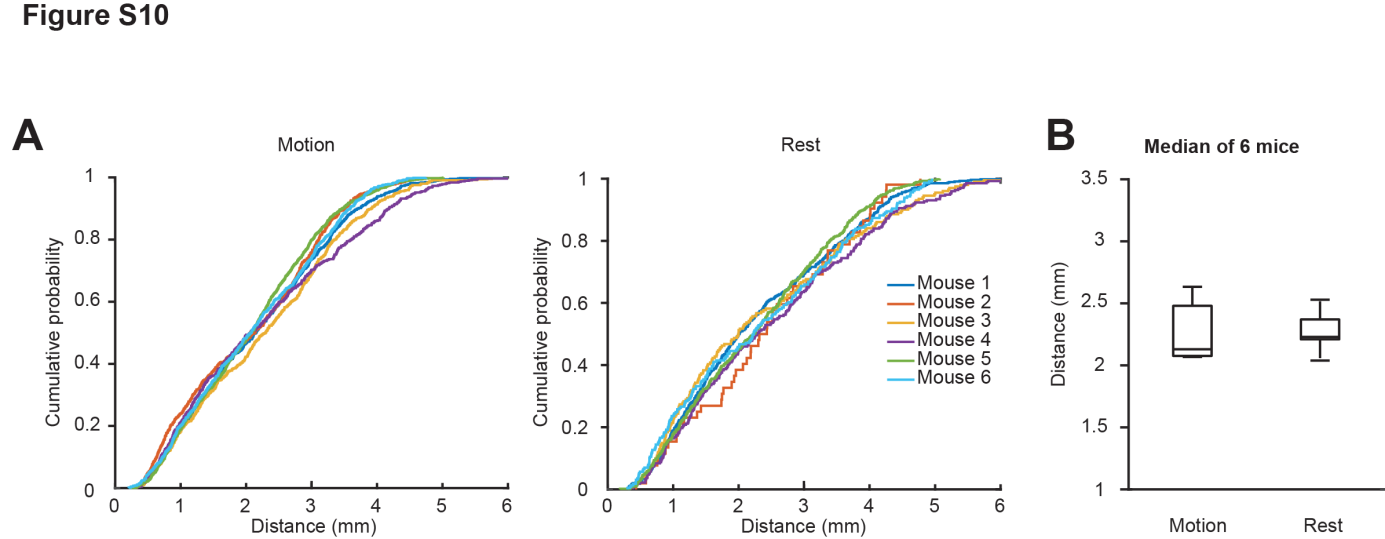

**Figure 5-2. Distance between simultaneously active cortical subregions.** ***A***, Cumulative probability functions of all pairwise distances between the centers of simultaneously active subregions belonging to the same multi-ROI filters during motion (left panel) and resting (right panel) time periods. Distributions obtained in different mice (n = 6) are shown in different colors. ***B***, Box-and-whisker plot showing median (per mouse) distances between the centers of coherently active subregions (n = 6 mice).

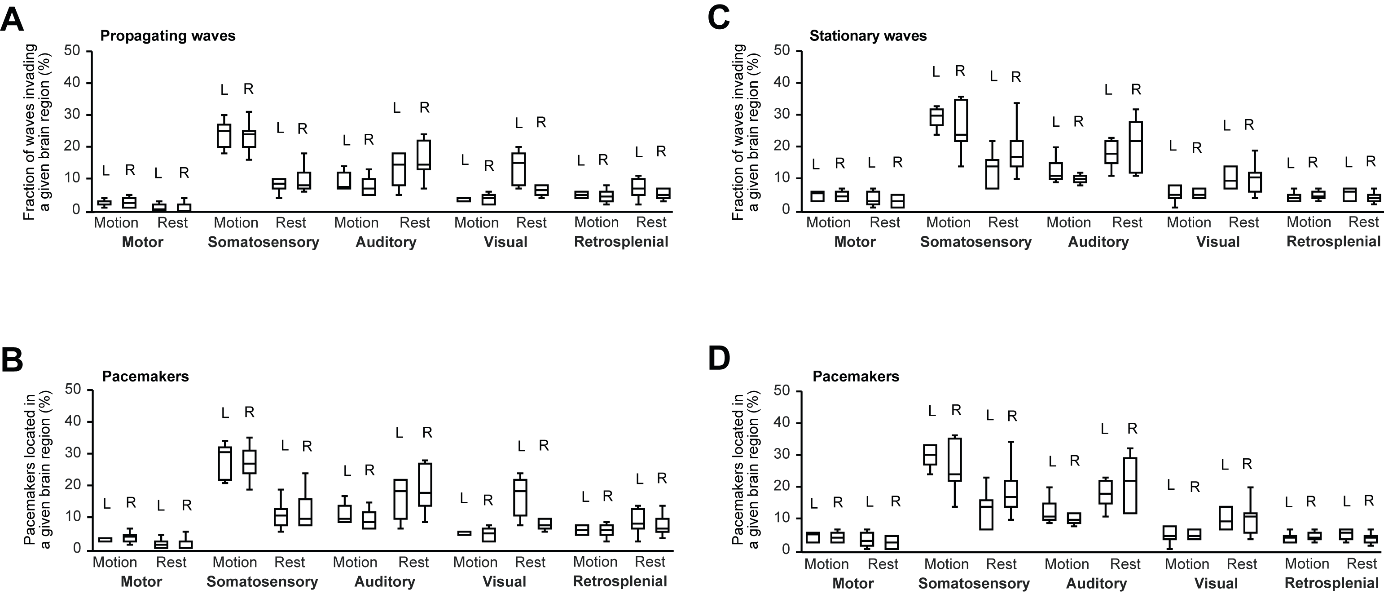

**Figure 5-3. Stationary and propagating waves in the neonatal mouse cortex. *A***, ***C***, Median (per mouse) fractions of propagating ***A***, and stationary ***B***, waves invading the respective cortical areas in the left (L) and the right (R) hemisphere. ***B****,* ***D***, Similar analyses as in ***A***, ***C*** but illustrating fractions of pacemakers for propagating ***B***, and stationary ***D***, waves located in the given cortical area (n = 6 mice).

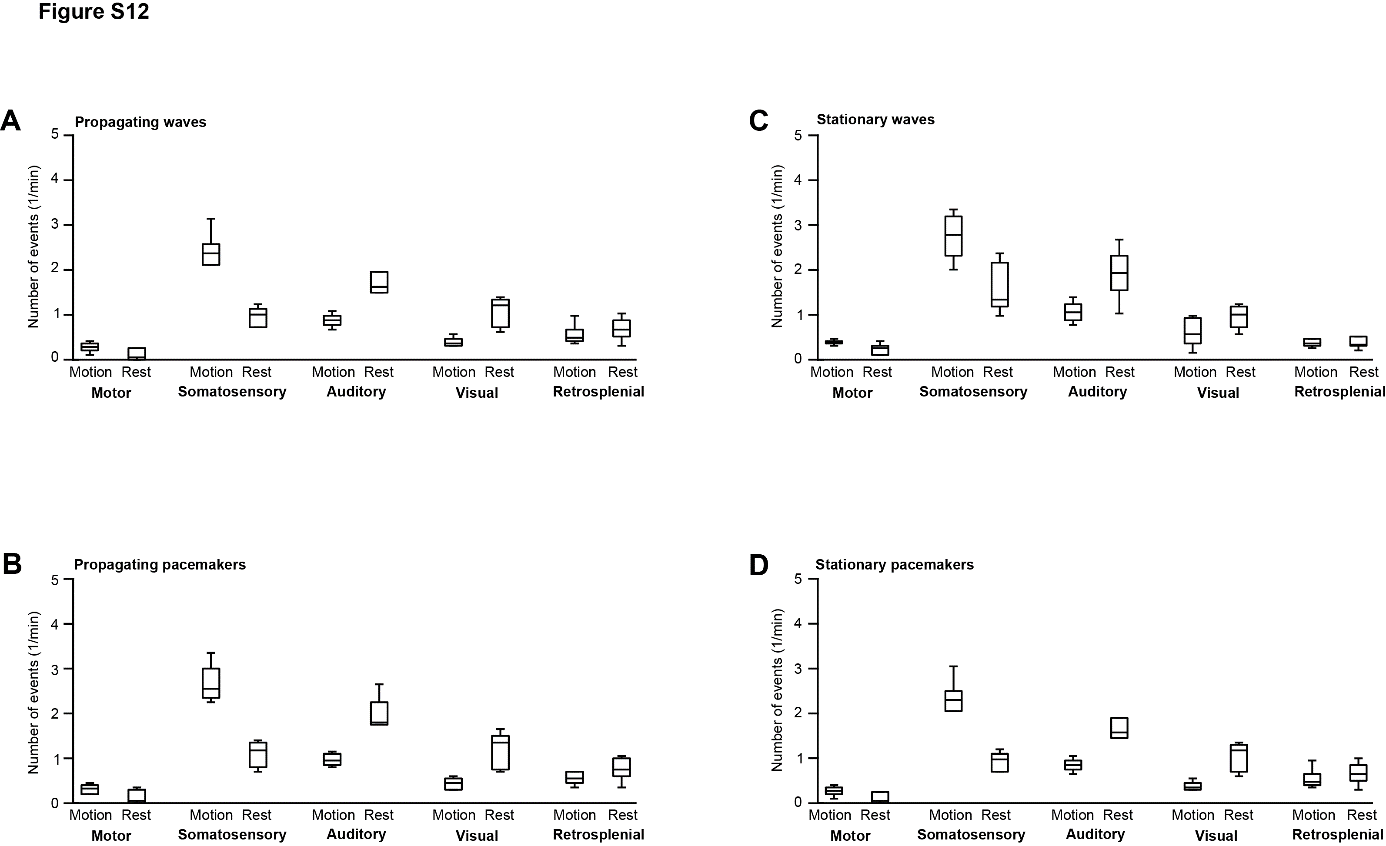

**Figure 5-4.** **The frequency of stationary and propagating Ca^2+^ waves in the neonatal mouse cortex.** ***A***, ***C***, Box-and-whisker plots show averaged between the two hemispheres median (per mouse) frequencies of propagating ***A***, and stationary ***C***, waves, invading the respective cortical regions. ***B***, ***D***, Similar analyses as in ***A***, ***C*** but illustrating median (per mouse) incidence of pacemaker events, located in the given cortical region, for propagating ***B***, and stationary ***D***, waves (n = 6 mice).

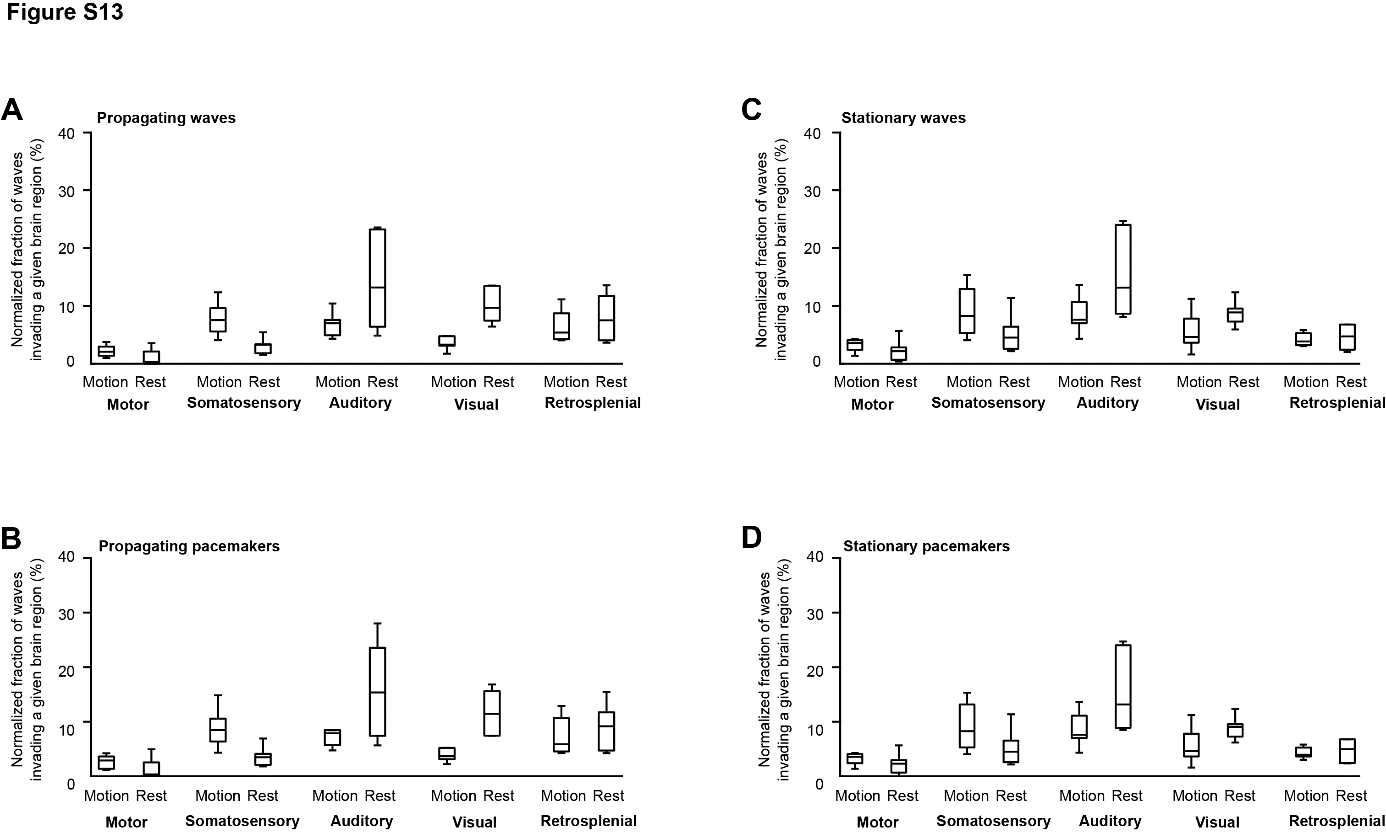

**Figure 5-5.** **Normalized fractions of stationary and propagating Ca^2+^ waves in the neonatal mouse cortex.** ***A***, ***C***, Box-and-whisker plots show averaged between the two hemispheres median (per mouse) normalized fractions of propagating ***A***, and stationary ***C***, waves, invading the respective cortical regions. ***B***, ***D***, Similar analyses as in ***A***, ***C*** but illustrating median (per mouse) normalized fractions of pacemakers for propagating ***B***, and stationary ***D***, waves located in the given cortical region (n = 6 mice). To calculate the normalized fractions, the respective region-specific values were first normalized to the area of the region and then normalized to the number of all propagating/stationary waves/pacemakers detected in this state.

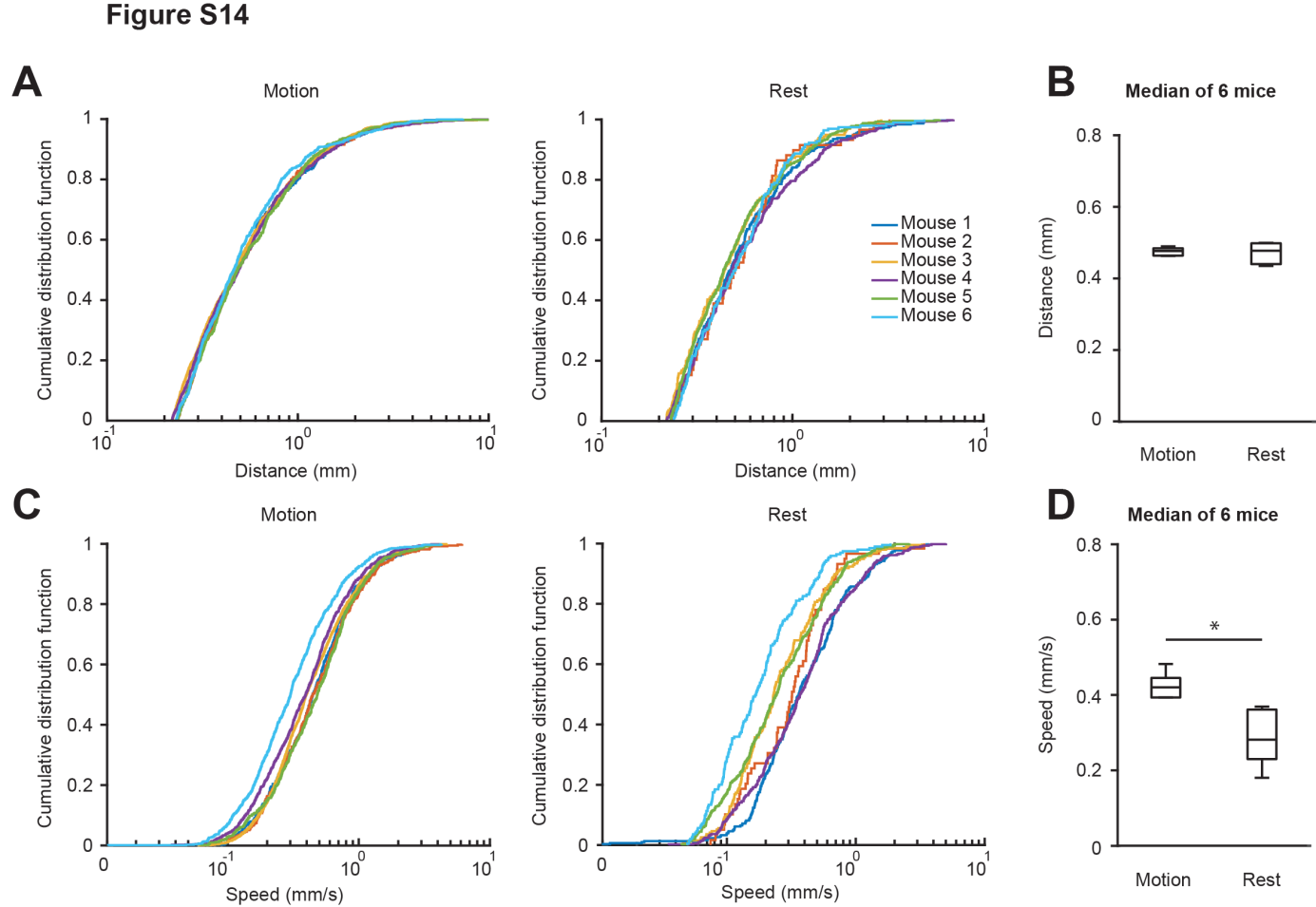

**Figure 5-6. Distance and speed of propagating waves.** ***A***, Cumulative probability functions of all distances traveled by propagating waves during motion (left panel) and resting (right panel) time periods. Distributions obtained in different mice (n = 6) are shown in different colors. X-axes have logarithmic scale. ***B***, Box-and-whisker plot showing median (per mouse) distances traveled by propagating waves during motion and resting time periods (paired Student’s t-test, t_5_ = 0.34, P = 0.75). ***C***, Cumulative probability functions of the average apparent speed of propagating waves recorded during motion (left panel) and resting (right panel) time periods. X-axes have logarithmic scale. ***D***, Box-and-whisker plot showing median (per mouse) apparent speed of propagating waves during motion and resting states. Obtained values are significantly different (paired Student’s t-test, t_5_ =4.3, P = 8 x 10^-3^).

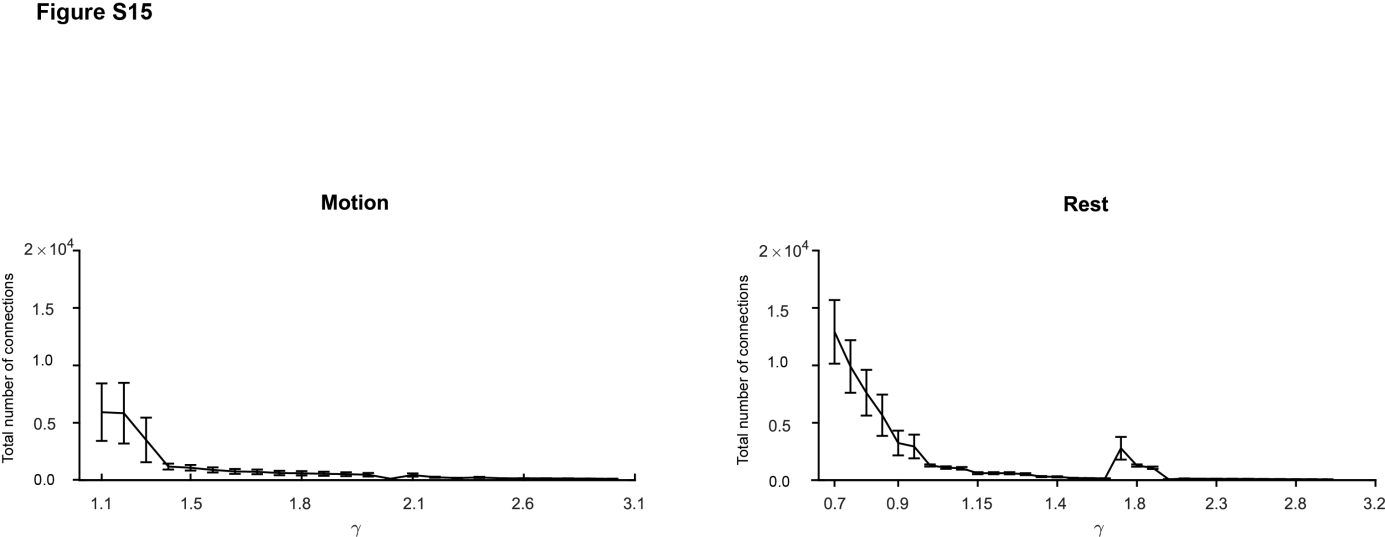

**Figure 6-1. Effect of the tuning parameter γ on the state-specific number of functional connections.** Total number of functional connections between the cortical regions during motion (left) and rest (right) as a function of γ. Data are shown as mean ± S.E.M (n = 6 mice).

**Movie 2-1. Patterns of endogenous activity in a wild type (C57BL/6) mouse.** Representative movie shows patterns of activity extracted using NMF algorithm. Left panel shows raw data. To make changes in intensity more visible, the mean (over time) value of each pixel was subtracted. Second left panel shows video, reconstructed using only blood vessel-containing NMF components. Second right panel shows video, reconstructed using only NMF components including large-scale (distributed) patterns. Right panel shows video, reconstructed using only NMF components containing local patterns of activity. Videos in all 4 panels are presented at 4x higher speed. The length of videos in real time is ~10 min. In each panel the intensity range is adjusted to maximize visibility. The white square in the upper right corner of the video marks the periods of animal’s movements.

**Movie 2-2. Patterns of endogenous activity in nestin-Cre x Ai95(RCL‐GCaMP6f)-D mouse.** Data are shown in the format described in Movie 2-1.
